# Simulation-based survey of TMEM16 family reveals that robust lipid scrambling requires an open groove

**DOI:** 10.1101/2024.09.25.615027

**Authors:** Christina A. Stephens, Niek van Hilten, Lisa Zheng, Michael Grabe

**Affiliations:** Cardiovascular Research Institute; Graduate Group in Biophysics; Department of Pharmaceutical Chemistry University of California, San Francisco, CA 94158

**Keywords:** TMEM16s, scramblases, membrane deformation, MD simulation

## Abstract

Biological membranes are complex and dynamic structures with different populations of lipids in their inner and outer leaflets. The Ca^2+^-activated TMEM16 family of membrane proteins plays an important role in collapsing this asymmetric lipid distribution by spontaneously, and bidirectionally, scrambling phospholipids between the two leaflets, which can initiate signaling and alter the physical properties of the membrane. While evidence shows that lipid scrambling can occur via an open hydrophilic pathway (“groove”) that spans the membrane, it remains unclear if all family members facilitate lipid movement in this manner. Here we present a comprehensive computational study of lipid scrambling by all TMEM16 members with experimentally solved structures. We performed coarse-grained molecular dynamics (MD) simulations of 27 structures from five different family members solved under activating and non-activating conditions, and we captured over 700 scrambling events in aggregate. This enabled us to directly compare scrambling rates, mechanisms, and protein-lipid interactions for fungal and mammalian TMEM16s, in both open (Ca^2+^-bound) and closed (Ca^2+^-free) conformations with statistical rigor. We show that all TMEM16 structures thin the membrane and that the majority of scrambling (>90%) occurs at the groove only when TM4 and TM6 have sufficiently separated. Surprisingly, we also observed 60 scrambling events that occurred outside the canonical groove, over 90% of which took place at the dimer-dimer interface in mammalian TMEM16s. This new site suggests an alternative mechanism for lipid scrambling in the absence of an open groove.

**Impact Statement:** The majority of TMEM16 lipid scrambling occurs in the open groove associated with Ca^2+^-activation, but limited scrambling also occurs in the dimer interface independent of Ca^2+^.

## Introduction

The TMEM16 family of eukaryotic membrane proteins, also known as anoctamins (ANO), is comprised of lipid scramblases (1–3), ion channels (4–7), and members that can facilitate both lipid and ion permeation (8–15). This functional divergence, despite their high sequence conservation, is a unique feature among the 10 vertebrate paralogues (16). So far, all characterized TMEM16s require Ca^2+^ to achieve their maximum transport activity, whether that be passive ion movement or lipid flow down their electrochemical gradients (8, 11, 17–21). TMEM16s play critical roles in a variety of physiological processes including blood coagulation (8, 22–24), bone mineralization (25), mucus secretion (26), smooth muscle contraction (27), and membrane fusion (28). Mutations of TMEM16 have also been implicated in several cancers (29–31), neuronal disorder SCAR10 (32, 33), and SARS-CoV2 infection (34). Despite their significant roles in human physiology, the functional properties of most vertebrate TMEM16 paralogues remain unknown. Moreover, even though we have significant functional and structural insight into the mechanisms of a handful of members (11, 13, 15, 35–58), it is still an open question whether all TMEM16s work in the same way to conduct ions or scramble lipids.

Over the past ten years, 63 experimental structures of TMEM16s have been determined, revealing a remarkable structural similarity between mammalian and fungal members despite the diversity in their functions. All structures, except for one of fungal *Aspergillus fumigatus* TMEM16 (afTMEM16) (59) which is a monomer, are homodimers with a butterfly-like fold (12, 15, 19, 21, 46, 49, 59–67), and each subunit is comprised of 10 transmembrane (TM) helices with the final helix (TM10) forming most of the dimer interface. Residues on TM6 form half of a highly conserved Ca^2+^-binding site that accommodates up to 2 ions. TM6 along with TM3, TM4, and TM5 also form a membrane spanning groove that contains hydrophilic residues that are shielded from the hydrophobic core of the bilayer in Ca^2+^-free states. When Ca^2+^ is bound, TM6 takes on a variety of conformational and secondary structural changes across the family, which can have profound effects on the shape of the membrane as seen in cryo-EM nanodiscs with TMEM16F (66). Ca^2+^-binding is also associated with the movement of the upper portion of TM4 away from TM6 which effectively exposes (opens) the hydrophilic groove to the bilayer, but this opening is not observed for all Ca^2+^ bound TMEM16 members (21, 46, 49, 61, 63–67).

It was first theorized (19, 68) and later predicted by molecular dynamics (MD) simulations (12, 35, 36, 40, 52, 55) that lipids can traverse the membrane bilayer by moving their headgroups along the water-filled hydrophilic groove (between TM4 and TM6) while their tails project into the hydrophobic center of the bilayer. This mechanism for scrambling, first proposed by Menon & Pomorski, is often referred to as the “credit card model” (69). All-atom MD (AAMD) simulations of open *Nectria haematococca* TMEM16 (nhTMEM16) have shown that lipids near the pore frequently interact with charged residues at the groove entrances (36), two of which are in the scrambling domain which confers scramblase activity to the ion channel-only member TMEM16A (70). Frequent headgroup interactions with residues lining the groove were also noted in atomistic simulations of open TMEM16K including two basic residues in the scrambling domain. Lipids experience a relatively low energy barrier for scrambling in open nhTMEM16 (<1 kcal/mol compared to 20-50 kcal/mol directly through the bilayer) (36, 69). Simulations also indicate that zwitterionic lipid headgroups stack in the open groove along their dipoles, which may help energetically stabilize them during scrambling (36, 42). Finally, simulations also show that lipids can directly gate nhTMEM16 groove opening and closing through interactions with their headgroups or tails (38, 39). It is important to note that all of these simulation observations are based on a limited number of spontaneous events from different groups (in aggregate we estimate that no more than 14 scrambling events have been reported in the absence of an applied voltage) (12, 36, 37, 40, 52, 55). Many more scrambling events (∼800 in aggregate) have been seen in coarse-grained MD (CGMD) simulations for nhTMEM16 (35), TMEM16K (12), mutant TMEM16F (F518H) and even TMEM16A (71); however, a detailed analysis of how scrambling occurred in these latter two was not provided. Moreover, a head-to-head comparison of fungal *versus* mammalian scrambling rates has not been made.

An outstanding question in the field is whether scrambling requires an open groove. This question has been triggered in part by the failure to determine wild type (WT) TMEM16F structures with open grooves wide enough to accommodate lipids, despite structures being solved under activating conditions (66, 67). Further uncertainty stems from data showing that scrambling can occur in the absence of Ca^2+^ when the groove is presumably closed (11–13, 19, 40, 43, 62). Moreover, afTMEM16 can scramble PEGylated lipids, which are too large for even the open groove (43). This last finding motivated Malvezzi *et al.* to propose an alternate model of scrambling (43) inspired by the realization that the bilayer adjacent to the protein, whether the groove is open or closed, is distorted in cryo-EM in nanodiscs (59–61, 66), MD simulations (12, 36, 38), and continuum models (36). The hallmark of this distortion is local bending and thinning adjacent to the groove (estimated to be 50-60% thinner than bulk for some family members) (12, 36, 38, 59–61, 66), and it has been suggested that this deformation, along with packing defects, may significantly lower the energy barrier for lipid crossing (36, 59, 66). To date no AAMD or CGMD simulation has reported scrambling by any wild-type TMEM16 harboring a closed groove; however, a CGMD simulation of the F518H TMEM16F mutant did report scrambling, but the details, such as whether the groove opened, were not provided (71). Again, since a comprehensive analysis across all family members has not been carried out, it is difficult to determine how membrane thinning is related to scrambling or if scrambling mechanisms are specific to certain family members, conformational states of the protein, or both. Additionally, lipids are also directly involved in how TMEM16 scramblases conduct ions. As first speculated in ref. (11), AAMD simulations have shown that ions permeate through the lipid headgroup-lined hydrophilic groove of TMEM16K and nhTMEM16 (37, 39, 41, 42, 58). How might this mechanism differ in the absence of an open groove?

To address these outstanding questions, we employed CGMD simulation to systematically quantify scrambling in 23 experimental and 4 computationally predicted TMEM16 proteins taken from each family member that has been structurally characterized: fungal scramblases nhTMEM16 and afTMEM16, and mammalian scramblase TMEM16K, TMEM16F, and TMEM16A (**Appendix 1-Table 1; Appendix 1-Figure 1**). CGMD, which was the first computational method to identify nhTMEM16 as a scramblase (35), enables us to reach much longer timescales, while retaining enough chemical detail to faithfully reproduce experimentally verified protein-lipid interactions (72). This allowed us to quantitatively compare the scrambling statistics and mechanisms of different WT and mutant TMEM16s in both open and closed states solved under different conditions (e.g., salt concentrations, lipid and detergent environments, in the presence of modulators or activators like PIP_2_ and Ca^2+^). Our simulations successfully reproduce experimentally determined membrane deformations seen in nanodiscs across both fungal and mammalian TMEM16s. They also show that only open scramblase structures have grooves fully lined by lipids and each of these structures promote scrambling in the groove with lipids experiencing a less than 1 kT free energy barrier as they move between leaflets. Interestingly, one simulation of TMEM16A, which is not a scramblase, initiated from a predicted ion conductive state scrambled lipids through a lipid-lined groove at a very low rate (only 2 events) suggesting that ion channel-only members may have residual non-detectable scramblase activity. Our analysis of the membrane deformation and groove conformation shows that most scrambling in the groove occurs when the membrane is thinned to a least 14 Å and the groove is open. We also observe 218 ion permeation events but only in well-hydrated systems with open grooves (98%) and a closed-groove TMEM16A structure (2%). Our simulations also reveal alternative scrambling pathways, which primarily occur at the dimer-dimer interface in mammalian structures.

**Figure 1.**
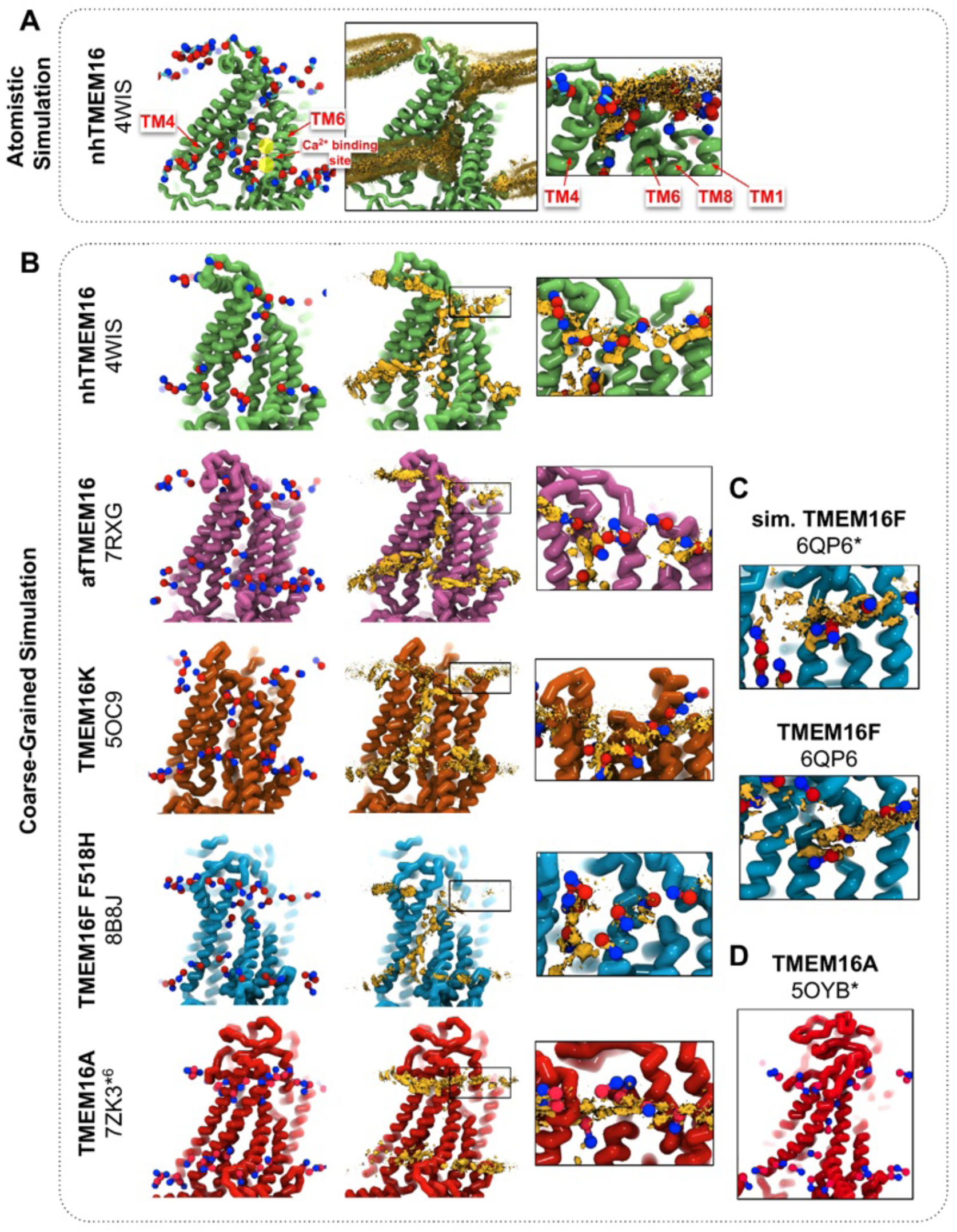
CG simulations of multiple TMEM16 structures capture lipid density in the TM4/TM6 pathway of scrambling competent members. **(A)** Snapshot and POPC headgroup density (right) from atomistic simulations of Ca^2+^-bound nhTMEM16 (PDB ID 4WIS) previously published in ref. (36). Only the lipid headgroup choline (blue) and phosphate (red) beads are shown for clarity. Density (brown isosurface) is averaged from both subunits across 8 independent simulations totaling ∼2 μs. Two yellow circles indicate the Ca^2+^-binding sites. **(B)** Snapshots from CG simulations of open Ca^2+^-bound nhTMEM16 (PDB ID 4WIS, green), afTMEM16 (PDB ID 7RXG, violet), TMEM16K (PDB ID 5OC9, orange), TMEM16F F518H (PDB ID 8B8J, blue), TMEM16K (orange), and TMEM16A (red). (C) Snapshots with lipid headgroup densities near simulated open (6QP6*) and closed (PDB ID 6QP6) TMEM16F. (D) Snapshot of simulated ion conductive TMEM16A (5OYB*). For each CG snapshot, again only the lipid headgroup choline (blue) and phosphate (red) beads are shown for clarity. Each density (brown isosurface) is averaged over both chains except TMEM16K and TMEM16A where only a single chain is used due to the structure’s asymmetry.

## Results

### Lipid densities from coarse-grained simulations match all-atom simulations and cryo-EM nanodiscs

We simulated Ca^2+^-bound and -free (apo) structures of TMEM16 proteins in a 1,2-dioleoyl-sn-glycero-3-phosphatidylcholine (DOPC) bilayer for 10 μs each using the Martini 3 force field (73). First, we determined how well the simulated membrane distortions matched experiment by comparing the annulus of lipids surrounding each protein to the lipid densities derived from structures solved in nanodiscs (**Figure 1-figure supplement 1**). The shapes of the membrane near the protein qualitatively match the experimental densities and the shapes produced from AAMD simulations and continuum membrane models (36, 59, 61). For example, the CG simulations capture the sinusoidal curve around both fungal scramblases in apo and Ca^2+^-bound states (**Figure 1-figure supplement 1**) previously determined by atomistic simulations (36) of Ca^2+^-bound nhTMEM16 (**Fig. 1A-B**). Even though membrane deformation is a general feature of TMEM16s, the shapes between fungal and mammalian members are noticeably different. Specifically, the membrane is flatter around TMEM16K and TMEM16F compared to the fungal members in both the nanodisc density and CGMD (**Figure 1-figure supplement 1**). For WT TMEM16s, whether the groove is open (**Fig. 1B***, insets*) or closed **(Figure 1-figure supplement 2),** strong lipid density exists near the extracellular groove entrances at TM1 and TM8. Interestingly, this density is lost in the simulation of the Ca^2+^-bound constitutively active TMEM16F F518H mutant (PDB ID 8B8J), consistent with what is seen in the cryo-EM structure solved in nanodisc (**Figure 1-figure supplement 1**). The lipid density is present, however, at this location for the simulated open Ca^2+^-bound WT TMEM16F (6QP6*, initiated from PDB ID 6QP6) and closed Ca^2+^-bound WT TMEM16F (PDB ID 6QP6) (**Fig. 1C**). The loss of density indicates that the normal membrane contact with the protein near the TMEM16F groove has been compromised in the mutant structure. Residues in this site on nhTMEM16 and TMEM16F also seem to play a role in scrambling but the mechanism by which they do so is unclear (59, 62, 66, 67).

Headgroup density isosurfaces from CGMD simulations of known scramblases bound to Ca^2+^ and with clear separation of TM4 and TM6 show that lipid headgroups occupy the full length of the groove creating a clear pathway that links the upper and lower membrane leaflets (nhTMEM16 (PDB ID 4WIS), afTMEM16 (PDB ID 7RXG), TMEM16K (PDB ID 5OC9) and TMEM16F F518H (PDB ID 8B8J), **Fig. 1B**). These simulation-derived densities crossing the bilayer are strikingly similar to lipids resolved in cryo-EM structures of fungal scramblases in nanodisc (59, 62). Individual simulation snapshots provide insight into how lipids traverse this pathway. Open-groove structures typically have an average of 2-3 lipids in the TM4/TM6 groove at any given time (**Figure 1-table supplement 1**). Additional analysis shows that all of the grooves are filled with water (**Appendix 1-Figure 2**). These profiles share additional features including a clear upward deflection of the membrane as it approaches TM3/TM4 from the left and a downward deflection as it approaches TM6/TM8 from the right; however, the degree of this deflection is not equal as can be seen for TMEM16K, which is less pronounced (**Fig. 1B**). These distortions are coupled to the sinusoidal curve around the entire protein, which was shown to thin the membrane across the groove and hypothesized to aid in scrambling (36).

**Figure 2.**
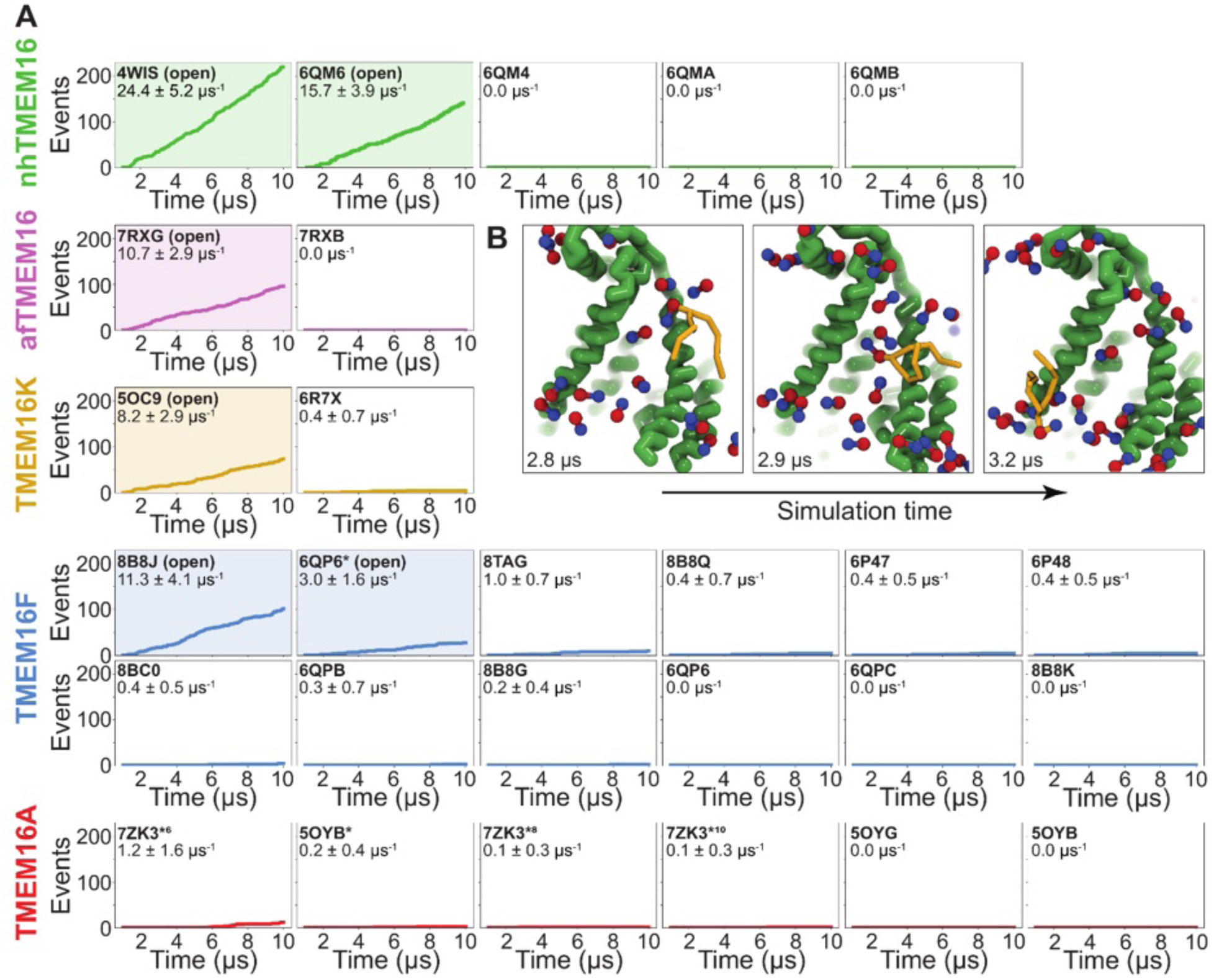
Simulated lipid scrambling differentiates closed/open conformations. **(A)** Accumulated scrambling events during CGMD simulations of experimental and simulated (sim) structures of nhTMEM16 (green), afTMEM16 (violet), TMEM16K (gold), TMEM16F (blue), and TMEM16A (red). Inset values are the average rate and its standard deviation. Plots corresponding to structures described as “open” in their original publications (PDB IDs 4WIS (19), 6QM6 (61), 7RXG (59), 5OC9 (12), 8B8J (60), and 6QP6* (55)) are shaded. **(B)** Snapshots of the open nhTMEM16 simulation (PDB ID 4WIS) showing a single scrambling event over time. The tail (yellow) of the scrambling lipid is explicitly shown, while all other lipids only show the phosphate (red)/choline (blue) headgroup.

Unlike the open Ca^2+^-bound scramblase structures, apo and closed Ca^2+^-bound TMEM16 structures lack lipid headgroup density spanning the bilayer, and their density profiles are more consistent across the entire family (**Figure 1-figure supplement 2**). The membrane is deformed near the groove with some lower leaflet lipid density entering part of groove and some of the upper leaflet density deflecting inward around TM1, TM6, and TM8 but not entering the closed outer portion of the groove. Again, the membrane around TMEM16F and TMEM16K is flatter than it is in the fungal scramblases. Similarly, simulation of a Ca^2+^-bound TMEM16A conformation that conducts Cl^-^ in AAMD (7ZK3*^6^, initiated from PDB ID 7ZK3, see Appendix 1-Methods and **Appendix 1-Figure 1**) samples partial lipid headgroup penetration into the extracellular vestibule formed by TM3/TM6, but lipids fail to traverse the bilayer as indicated by the lack of density in the center of the membrane (**Fig. 1B**). This finding is consistent with TMEM16A lacking scramblase activity (54, 70); however, we simulated another ion-conductive TMEM16A conformation that can achieve a fully lipid-lined groove during its simulation (**Fig. 1D**), although this configuration was uncommon (**Figure 1-table supplement 1**).

### Simulations recapitulate scrambling competence of open and closed structures

To quantify the scrambling competence of each simulated TMEM16 structure, we determined the number of events in which lipids transitioned from one leaflet to the other (see Methods, **Figure 2-figure supplement 1**). The scrambling rates calculated from our MD trajectories are in excellent agreement with the presumed scrambling competence of each experimental structure (**Fig. 2A**). The strongest scrambler was the open-groove, Ca^2+^-bound fungal nhTMEM16 (PDB ID 4WIS), with 24.4 ± 5.2 events per μs (**Figure 2-figure supplement 2A**). In line with experimental findings, the open Ca^2+^-free structure (PDB ID 6QM6), which is structurally very similar to PDB ID 4WIS, also scrambled lipids (15.7 ± 3.9 events per μs, **Figure 2-figure supplement 2B**) (61). In contrast, we observed no scrambling events for the intermediate-(PDB ID 6QMA) and closed-(PDB ID 6QM4, PDB ID 6QMB) groove nhTMEM16 structures. We observed a similar trend for the fungal afTMEM16, where our simulations identified the open Ca^2+^-bound cryo-EM structure (PDB ID 7RXG) as scrambling competent (10.7 ± 2.9 events per μs, **Figure 2-figure supplement 2C**) while the Ca^2+^-free closed-groove structure (PDB ID 7RXB) was not.

For TMEM16K, our simulations showed that the Ca^2+^-bound X-ray structure (PDB ID 5OC9) facilitates scrambling (8.2 ± 2.9 events per μs) in line with experiments in the presence of Ca^2+^, when the groove is presumably open, and previous MD simulations (12). Interestingly, we found a stark asymmetry in the number of scrambling events between the two monomers, with >80% of events happening via chain B (**Figure 2-figure supplement 3A**). Although both monomers are Ca^2+^-bound, chain B has a slightly wider ER lumen-facing entrance to the groove in the experimental starting structure (PDB ID 5OC9, **Figure 2-figure supplement 3B**) and spontaneously opened its groove more than subunit A during the simulation (8.2 ± 1.3 Å compared to 5.8 ± 0.6 Å on average, **Figure 3-figure supplement 6A**), which likely accounts for the increased rate. The closed-groove TMEM16K conformation (PDB ID 6R7X) showed very little scrambling activity (0.4 ± 0.7 events per μs).

**Figure 3.**
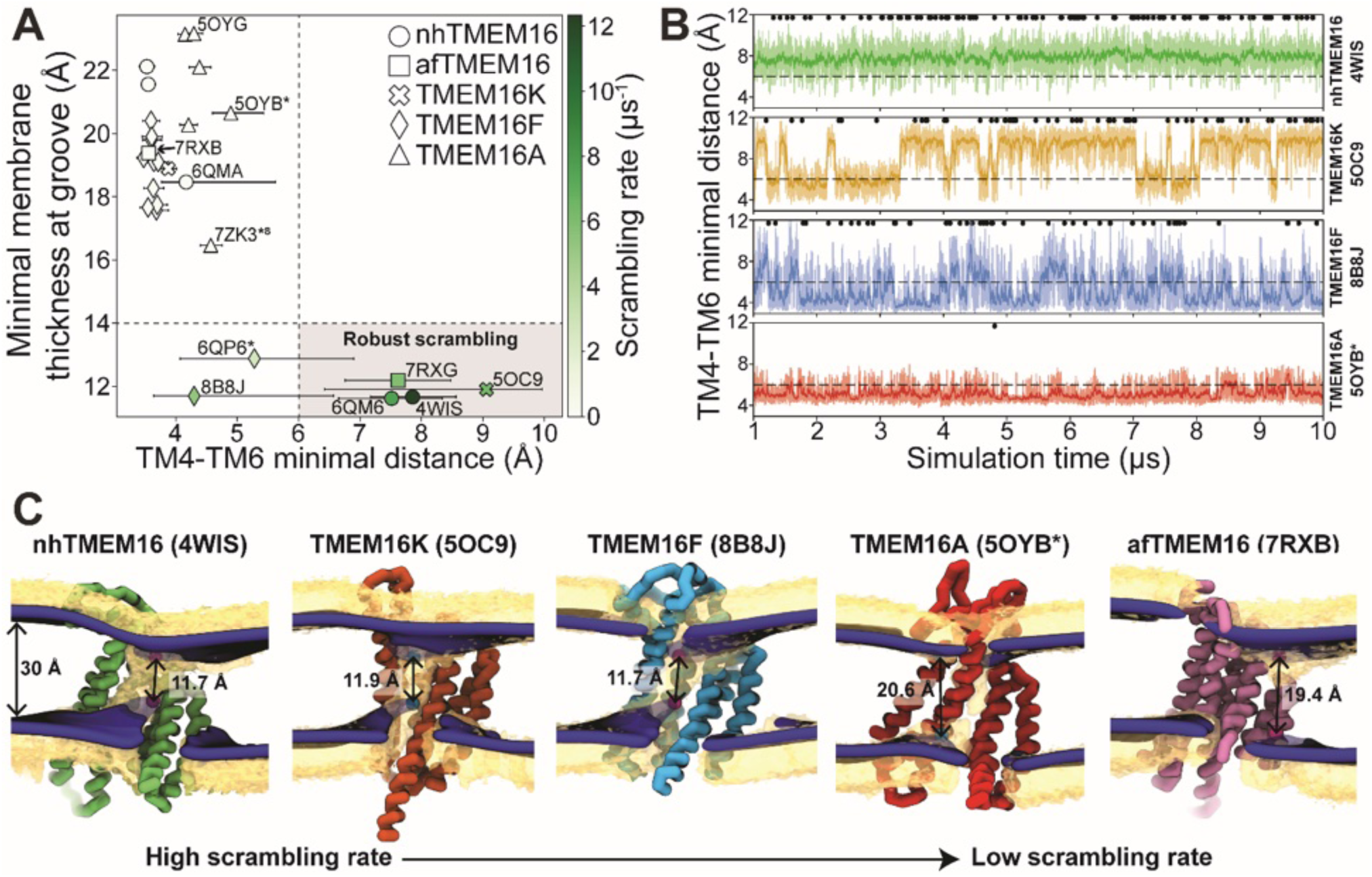
Lipid scrambling rates correlate with groove openness and membrane thinning. **(A)** The minimal membrane thickness at the groove plotted against the median width of the groove measured based on the minimal distance between any two residues on TM4 and TM6 of the groove with the most scrambling events. The lower and upper error bars represent the 25% (Q1) and 75% (Q3) quartiles, respectively. Each data point is colored by the scrambling rate through that same groove. Dashed lines define minimal TM4-TM6 distance and membrane thickness requirements for robust scrambling (shaded grey quadrant). **(B)** Simulation time traces of the TM4-TM6 minimal distance at the most scrambling-competent groove of 4WIS, 5OC9, 8B8J, and 5OYB* (top to bottom). The dashed line indicates the 6 Å threshold we defined for scrambling-competent groove opening. Black dots indicate time points at which a scrambling event is completed. The solid curve is a recursively exponentially weighted moving average with smoothing factor 0.1, while the transparent curve is the raw distance values. **(C)** Density isosurfaces for DOPC headgroup beads (yellow) and average membrane surface calculated from the glycerol beads (blue) for representative nhTMEM16, TMEM16K, TMEM16F, TMEM16A, and afTMEM16 simulations. Panels are ordered left to right by decreasing scrambling rate. Cartoon beads and arrows in each image indicate the closest points between the inner and outer leaflet of the average surface.

Although TMEM16F is a known lipid scramblase found in the plasma membrane of platelets (10), none of the WT structures solved to date, even those determined under activating conditions, have exhibited an open hydrophilic groove. We simulated 10 of these proteins and observed little to no lipid scrambling in each case (**Fig. 2*A***). Others have shown that mutations at position F518 turns TMEM16F into a constitutively active scramblase (52). The F518H mutant (PDB ID 8B8J) is structurally characterized by a kink in TM3, and TM4 pulls away from TM6 35° compared to a closed WT TMEM16F structure (PDB ID 6QP6) (60). In our simulations, TMEM16F F518H (PDB ID 8B8J) was the only system initiated directly from a solved structure that showed scrambling activity (11.3 ± 1.6 events per μs, **Figure 2-figure supplement 4A**). Additionally, we performed CGMD on a WT TMEM16F with a single open groove obtained from AAMD initiated from a closed-state structure (cluster 10 in (55), 6QP6* in **Fig. 2A**). We observed moderate lipid scrambling activity (3.0 ± 1.6 events per μs), most of which happened through the open groove (**Figure 2-figure supplement 4B, C**). Although the rates of scrambling are higher for the mutant than the open WT TMEM16F, there were no noticeable differences in how lipids enter the pathway or how long they take to transition (**Figure 4-figure supplement 3**).

**Figure 4.**
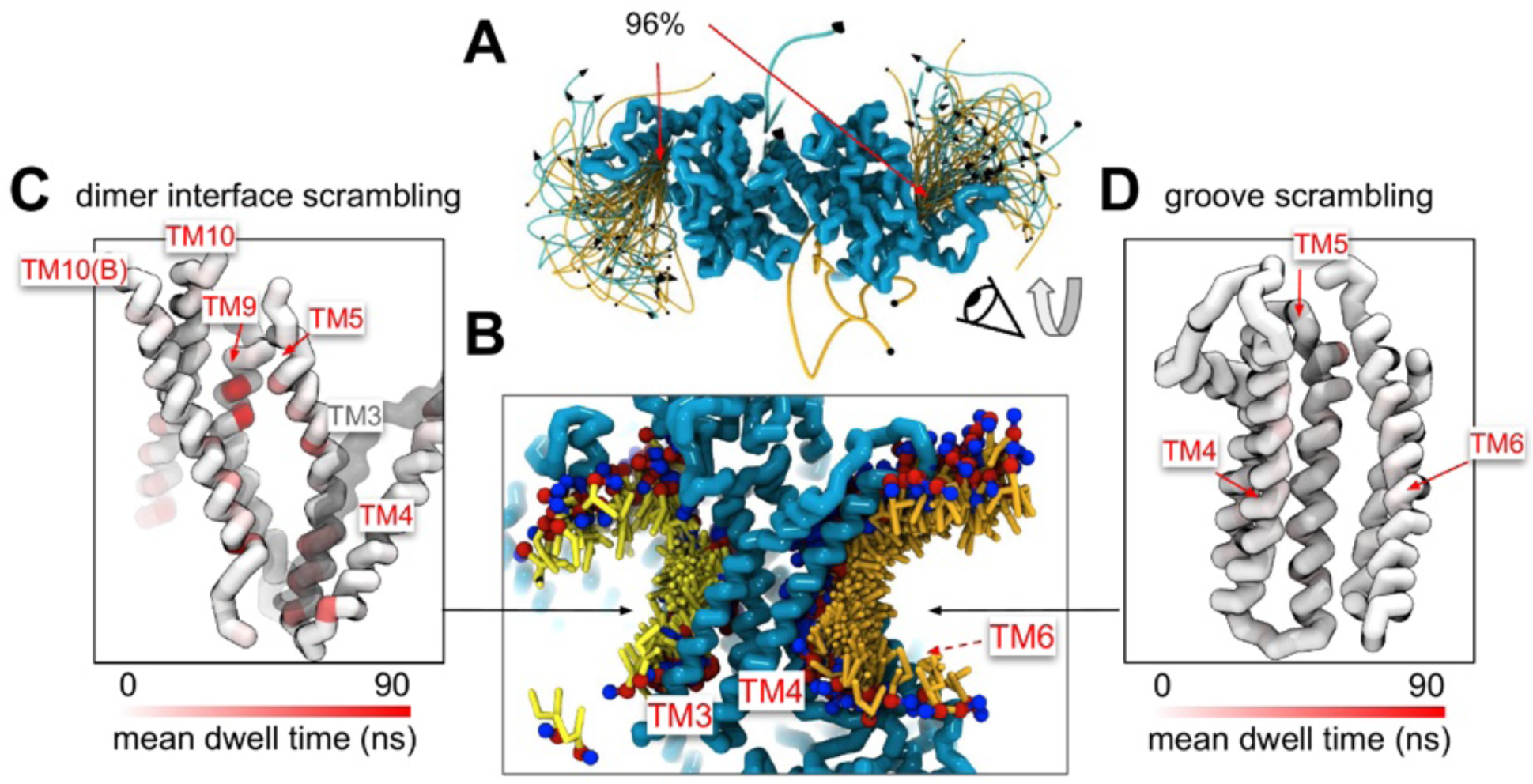
Lipid scrambling events and lipid-protein residue contact in the dimer interface and canonical TM4/TM6 groove. **(A)** Traces of all scrambling lipids in a TMEM16F (PDB ID 8B8J) simulation. Lipid scrambling from the inner to outer leaflet are illustrated as cyan traces and from the outer to inner leaflet as yellow traces. **(B)** Cartoon depiction of two individual inward scrambling events along the TM4/TM6 groove (orange tail with red/blue headgroup) and the dimer interface (yellow tail with red/blue headgroup) with multiple snapshots over time. Only headgroup, first and second tail beads are shown for clarity. **(C,D)** Protein backbone colored by mean lipid headgroup interaction (dwell) time at the TMEM16F dimer interface (C) and TM4/TM6 groove (D).

Finally, we simulated six structures of mouse TMEM16A, which functions as an ion channel but lacks lipid scrambling activity (46). As expected, both the Ca^2+^-bound (PDB ID 5OYB) and the Ca^2+^-free (PDB ID 5OYG) experimental structures failed to induce scrambling in the CGMD simulations, as did one alternative and two ion conduction-competent structures that were obtained from AAMD (see Appendix 1-Methods for details). However, a TMEM16A state with an open hydrophilic groove predicted by Jia & Chen (5OYB*, simulations initiated from PDB ID 5OYB (48)) did scramble a single lipid through each groove in a manner nearly identical to the scramblases (**Figure 2-figure supplement 5A, E** and **Figure 3-video 4).**

### Groove dilation is the main determinant for scrambling activity

The relative impact of membrane thinning versus TM4/TM6 groove opening on the lipid scrambling rate has long been debated in the TMEM16 field. One of the primary open questions is whether membrane thinning is *sufficient* for scrambling when the groove is closed (74). In our CGMD simulations, 92% of the observed scrambling events occur along TM4 and TM6 with headgroups embedded in the open hydrated groove, in line with the credit card model, which we refer to as “in-the-groove” scrambling (**Table 1**). To visualize how groove openness and membrane thinning relate to these events, we plotted the minimum distance between residues on TM4 and TM6 against the minimal thickness near the groove in our average membrane surfaces (see Methods for details) and colored each data point by scrambling rate in the groove (**Fig. 3A**).

**Table 1.** Number of scrambling events in and out of the canonical groove pathway. Scrambling events where the lipid headgroup transitions between leaflets within 4.7 Å of the DOPC maximum density pathway. All other events were considered “out-of-the-groove”. For the full list of simulations and scrambling rates see source data file.

| homolog | PDB code | # of in-the-groove events | # of out-of-the-groove events | total | average scrambling rate ($\mu\text{s}^{-1}$) |
| --- | --- | --- | --- | --- | --- |
| nhTMEM16 | 4WIS | 219 | 1 | 220 | 24.4±5.2 |
| nhTMEM16 | 6QM6 | 141 | 0 | 141 | 15.7±3.9 |
| afTMEM16 | 7RXG | 96 | 0 | 96 | 10.7±2.9 |
| TMEM16K | 5OC9 | 66 | 8 | 74 | 8.2±2.9 |
| TMEM16K | 6R7X | 0 | 4 | 4 | 0.4±0.7 |
| TMEM16F F518H | 8B8J | 98 | 4 | 102 | 11.3±4.1 |
| TMEM16F | 6QP6* | 24 | 3 | 27 | 3.0±1.6 |
| TMEM16F T137Y | 8TAG | 0 | 9 | 9 | 1.0±0.7 |
| TMEM16F | 6P47 | 1 | 3 | 4 | 0.4±0.5 |
| TMEM16F | 6P48 | 0 | 4 | 4 | 0.4±0.5 |
| TMEM16F F518H/Q623A | 8BC0 | 0 | 4 | 4 | 0.4±0.5 |
| TMEM16F F518H | 8B8Q | 0 | 4 | 4 | 0.4±0.7 |
| TMEM16F F518H | 8B8G | 2 | 0 | 2 | 0.2±0.4 |
| TMEM16F | 6QPB | 0 | 3 | 3 | 0.3±0.7 |
| TMEM16A | 7ZK3*<sup>6</sup> | 0 | 11 | 11 | 1.2±1.6 |
| TMEM16A | 5OYB* | 2 | 0 | 2 | 0.2±0.4 |
| TMEM16A | 7ZK3*<sup>10</sup> | 0 | 1 | 1 | 0.1±0.3 |
| TMEM16A | 7ZK3*<sup>8</sup> | 0 | 1 | 1 | 0.1±0.3 |

Interestingly, *all* the TMEM16 structures included in this study thin the membrane to 23 Å or less, which is at least 7 Å thinner than the bulk membrane thickness (30 Å), regardless of scrambling activity **(Fig. 3A**, **Figure 3-figure supplement 1-5)**. We observed negligible scrambling activity (0-1 events in the groove) in grooves that fail to thin the membrane less than 14 Å and at the same time do not or very rarely sample TM4-TM6 distances above 6 Å (**Fig. 3A**, upper left quadrant). On the other hand, all active scramblers have a minimal bilayer thickness below 14 Å. Among these structures, we observed the highest scrambling rates in grooves that remain open, with TM4-TM6 distances above 6 Å, throughout most of the simulation (shaded region). To the left of this shaded area there are two TMEM16F structures (PDB ID 8B8J and 6QP6*) that spent less than half of their simulation time in an open configuration (note large error bars, and **Fig. 3B**) and had scrambling rates similar to (PDB ID 8B8J) or less than half of (6QP6*) rates for the open scramblases (PDB IDs 6QM6, 7RXG, and 5OC9). Although these results indicate that scrambling rates are generally higher with thinner membranes and wider grooves, we want to clarify that lipids flowing into the upper and lower vestibules of the dilated grooves heavily contribute the observed <14 Å membrane thickness (**Fig. 3C**). Therefore, we argue that the extremely thin membranes are likely correlated with groove opening, rather than being an independent contributing factor to lipid scrambling. Thus, the major determinant of lipid scrambling by TMEM16s is dilation of the TM4/TM6 groove. Upon closer inspection of TMEM16F, we noticed that hydrophobic residues (H/F518, W619, and M522) at the midpoint of the pathway, previously identified as an activation gate (52), dynamically swing open to sporadically allow lipids through (**Figure 3-figure supplement 6C; Figure 3-video 3 and 5)**. Although the distribution of the groove distances is similar for both TMEM16F structures that exhibit scrambling (**Figure 3-figure supplement 6B**), the WT open structure (6QP6*) has half the single subunit scrambling rate. We observed similar fluctuations in both subunits of the open asymmetric TMEM16K (PDB ID 5OC9) which transiently constrict the lipid pathway at Y366/I370/T435/L436 (**Figure 2-figure supplement 3A; Figure 3-figure supplement 6A and D**; **Figure 3-video 2 and 6**). Again, we observed that the subunit with more scrambling activity (8 times more) spent more time in an open groove configuration (**Figure 3-figure supplement 6B**). Time traces of the TM4-TM6 distances emphasize the two-state, discrete nature of the TMEM16K groove as it opens and closes, the consistently open nature of nhTMEM16 with small fluctuations, and then the frequent fluctuations of the TMEM16F F518H mutant, which has a running average that constantly flickers from 4 Å to the 7-8 Å range (**Fig. 3B**). Qualitatively, scrambling occurs more frequently when the groove is open for TMEM16K and F (black dots in **Fig. 3B**), while the consistently open Ca^2+^-bound nhTMEM16 structure (PDB ID 4WIS) allows lipid headgroups to scramble in an uninterrupted fashion (**Fig. 3B**, **Figure 3-video 1**).

Although all structures of TMEM16A, which is not a scramblase, have negligible scrambling in the groove, we did observe two events for a predicted ion conductive state (5OYB*) which samples an average TM4-TM6 distance very close, but just below, to the empirically determined 6 Å threshold for scrambling (**Fig. 3B**). We observed lipid headgroups throughout the pathway, but just as for the TMEM16F and K structures, the flow of lipid is obstructed by residues at the center of the groove (I550, I551, and K645), and in TMEM16A they more rarely separate to allow lipids to pass (**Figure 3-figure supplement 6B**). Lipids are also notably more stagnant in the pore than in the open TMEM16Fs and appear to be stabilized by electrostatic interactions with two charged residues, E633 and K645 (**Figure 3-video 4**).

Despite these individual differences in groove dynamics, scrambling occurs in an identical manner across the family. Scrambling lipids move through the TM4/TM6 groove quickly, with dwell times for individual lipids below 20 ns. However, we observed longer dwell times for TMEM16K and TMEM16F at the groove constriction points whereas in other scramblases the dwell times are more evenly distributed along the groove (**Figure 4-figure supplement 3-4**). Among the scramblases, the free energy profile for lipids moving through the open groove is barrierless (<1 kT) (**Appendix 1-Figure 3A**) with similar kinetics among the homologs and a mean diffusion coefficient between 10 and 16 Å^2^/ns (**Appendix 1-Figure 4**). Scrambling events also enter and leave the groove at random locations (**Figure 2-figure supplement 2-5**) with only 3-10% of events passing through the high-density lipid regions on lower TM4 and upper TM6/TM8 (**Fig. 1B, Figure 4-figure supplement 2**). We previously identified four residues (E313, R432, K353, and E352) at the intracellular and extracellular entrances of the nhTMEM16 groove that we hypothesized help organize or stabilize scrambling lipids ((36), **Figure 3-figure supplement 1A-B**). However, our CGMD of the same nhTMEM16 structure shows that although these residues have elevated contact frequencies, more than half of the contacts are made with bulk lipids that never scramble (**Figure 4-figure supplement 1C, D**). Lastly, the in-the-groove scrambling events were Poisson distributed for all open and transiently open scramblases (**Appendix 1-Figure 5**), indicating lipids do not scramble in a regular or kinetically coordinated fashion.

### Water and ion content in the groove

To quantify how hydration of the groove or pore relates to scrambling, we measured the number of water permeation events along the pathway of maximum water density at the grooves (**Appendix 1-Figure 2A**; all values in source data file; see Appendix 1-Methods for details). As expected, permeation through the closed scramblase structures was low, < 30 events per μs on average, while dilated TM4/TM6 grooves (5 out of 6 Ca^2+^-bound) support 300-550 permeation events per μs on average. Nonetheless, even when the groove is inaccessible to lipids in closed and intermediate states, including the TMEM16A ion channel path, it remains hydrated with the waters shielded from the hydrophobic core of the membrane (**Appendix 1-Figure 2A***, closed*). As the groove opens, water is exposed to the membrane core and lipid headgroups insert themselves into the water-filled groove to bridge the leaflets (**Appendix 1-Figure 2A,** *open*) as observed in fully atomistic simulations (12, 35–37, 39–42, 52, 55, 58).

We also observed spontaneous permeation of Na^+^ and Cl^-^ ions through the scramblase TMEM16 grooves and TMEM16A pore (**Appendix 1-Figure 2B**; number of permeation events in source data file), in line with the known ion-conducting capacity of these proteins (8–15, 41, 42). Of the fungal structures, only the scrambling competent open states sampled multiple ion permeation events with Ca^2+^-bound nhTMEM16 (PDB ID 4WIS) showing highest conductance followed by Ca^2+^-free nhTMEM16 (PDB ID 6QM6), which was 3 times lower, and then Ca^2+^-bound afTMEM16 (PDB ID 7RXG), which was another 3 times lower again. We also measured cation-to-anion selectivity ratios of 5.1, 3.2, and 6 for each simulation, respectively, computed from the ratio of total counts (P*_Na_*/P*_Cl_*). Our simulations are consistent with experiments showing that both fungal scramblases transport anions and cations (13), and both are weakly cation selective (P*_K_*/P*_Cl_* = 1.5 for afTMEM16 based on experiment (11) and P*_Na_*/P*_Cl_* = 8.7 for nhTMEM16 based on AAMD (42)). Our CGMD simulations also sample ion conduction through open Ca^2+^-bound TMEM16F F518H (PDB ID 8B8J, P*_Na_*/P*_Cl_* = 1.3), which had the most ion permeation events (102) across the family, simulated open TMEM16F (6QP6*, P*_Na_*/P*_Cl_* = 0.33), and open TMEM16K (PDB ID 5OC9, P*_Na_*/P*_Cl_* = 1.8). This latter result on TMEM16K qualitatively agrees with experiment showing a slight cation preference (12), while experimental results for TMEM16F are more complex as its ion selectivity depends on membrane potential and divalent/monovalent cation concentrations (75–77). Our simulation of the TMEM16F F518H mutant in 150 mM NaCl is most close to whole cell recordings performed in intracellular 150 mM NaCl and 15 μM Ca^2+^ where P*_Na_*/P*_Cl_* = 1.0 ± 0.1 (77), which is very similar to our simulated value of 1.3. With regard to the selectivity values reported here, it is important to note that we observed less than 20 total events each for WT TMEM16F (6QP6*), afTMEM16, and TMEM16K (see source data file), and therefore, the values are prone to statistical error. We are more confident in the ratios reported for TMEM16F F518H and nhTMEM16 (PDB ID 4WIS) as those emitted 99 and 61 events, respectively.

Finally, TMEM16A (7ZK3*^8^) had 4 Cl^-^ and no Na^+^ permeation events, consistent with its experimentally measured anion selectivity (P*_Na_*/P*_Cl_* = 0.1 (45)). Interestingly, we did not observe Cl^-^ permeation in any of the other computationally predicted TMEM16A structures (5OYB*, 7ZK3*^8^, and 7ZK3*^10^), while AAMD simulations of these structures all reported Cl^-^ conduction (48).

### Scrambling also occurs out-of-the-groove

A minority of our observed scrambling events (8%) occurred outside of the hydrophilic groove between TM4 and TM6. Surprisingly, most of these events happened at the dimer interface with lipids inserting their headgroups into the cavity outlined by TM3 and TM10 (**Fig. 4A-B**; **Figure 2-figure supplement 2-5**). We only observed scrambling at this location in simulations of the mammalian homologs. In atomistic simulations of a closed Ca^2+^-bound TMEM16F (PDB ID 6QP6), we observed a similar flipping event for a POPC lipid into the dimer interface (**Figure 4-figure supplement 5)**. Although the dimer interface is largely hydrophobic, there are a few polar and charged residues in the cavity near the membrane core and water is present in the lower half of the cavity (**Fig. S24**). In fact, the headgroup of the lipid in our atomistic simulation of TMEM16F interacts with a glutamate (E843) and lysine (K850) on TM10 near the membrane midplane (**Figure 4-figure supplement 5**). Lipids that scramble at the dimer interface interact with the protein up to 10-fold longer on average than those in the canonical groove (**Fig. 4C-D**). The most prolonged interactions occur at sites containing aromatic residues into which the lipid tails intercalate (**Figure 4-figure supplement 7**).

There were five more out-of-the-groove events including one that occurred across a closed TM4/TM6 groove of Ca^2+^-bound TMEM16F (PDB ID 6P47). From all our observed scrambling events, this is the only one that fits the postulated out-of-the-groove definition where scrambling is expected to take place near TM4/TM6 but without inserting into the groove (78) (**Appendix 1-Figure 6A**). Two events occurred concurrently along TM6 and TM8 again near the hydrophilic groove of a Ca^2+^-bound closed TMEM16F (PDB ID 8TAG) (**Appendix 1-Figure 6A-B**). Lastly, two events occurred along TM3 and TM4, one near the canonical TM4/TM6 groove of an open nhTMEM16 (PDB ID 4WIS) and the other adjacent to the pore of an ion conductive TMEM16A (7ZK3*^8^) (**Appendix 1-Figure 6C-D**). In each of these five out-of-the-groove events, the scrambling lipid traverses with 2-4 water molecules around its headgroup.

## Discussion

Previous all-atom simulations of TMEM16 have captured partial translocations or – at most – a handful of complete scrambling events (e.g., (36, 40, 55)) due to the challenges inherent in simulating molecular events on the low microseconds time scale. Although these small number of AAMD-derived scrambling events yielded key insights into specific protein-lipid interactions and scrambling pathways, they cannot provide rigorous statistics on scrambling rates, nor can they be leveraged to perform a large high-throughput comparison between the various family members. To circumvent sampling issues, we used CGMD to systematically quantify lipid scrambling by five TMEM16 family members and relate their scrambling competence to their structural characteristics and ability to distort the membrane. Our simulations correctly differentiate between open and closed conformations across the five family members, consistent with a recent study that showed good qualitative agreement between *in vitro* and *in silico* lipid scrambling using the same Martini 3 force field on a diverse set of proteins, including some TMEM16s (79). In addition to lipid scrambling ability, our results are in accord with the general finding that TMEM16s show very little to no ion selectivity, although permeability ratios vary depending on ion concentrations and lipid environments (42, 75–77). Because the simulation conditions and system setups were identical in all our simulations, we are in a unique position to directly compare a host of biophysical properties between different TMEM16 family members and their structures to answer ongoing questions in the field.

In our simulations, *all* TMEM16 structures thin the membrane by at least 7 Å, while some pinch the membrane by as much as 18 Å resulting in leaflet-to-leaflet distances at the groove of just 12 Å (**Fig. 3*A, D***). We (36) and others (12, 39, 59, 60, 66) have hypothesized that thinning lowers the physical and energetic barrier for lipid scrambling, but what is surprising is that even non-scrambling, closed-groove structures elicit such large membrane distortions. For instance, several of the closed groove TMEM16F structures and the ion channel TMEM16A (7ZK3^*8^) compress the membrane 13-14 Å. Despite this large deformation, these conformations do not induce scrambling. On the other hand, structures with dilated grooves exhibit robust scrambling and thin the membrane another 3-4 Å, resulting in the most distorted bilayers. However, because this extreme membrane thinning is coupled to lipid entry into the upper and lower vestibules upon groove opening, it is difficult to determine how much the membrane thinning alone contributes to the resulting scramblase activity. Thus, we conclude that groove dilation is the ultimate trigger for rapid lipid scrambling, and the importance of membrane thinning to modulating scrambling rates has yet to be determined.

Of the scrambling competent TMEM16 structures, the open groove nhTMEM16 (PDB ID 4WIS) is the fastest scrambler, with a rate twice as high as the other homologs (**Fig. 2A** and **3A-B**). Yet on average its groove width and membrane thinning are similar (within 1-2 Å) to the other robust scramblers nhTMEM16 (PDB ID 6QM6), afTMEM16 (PDB ID 7RXG), and TMEM16K (PDB ID 5OC9) (**Fig. 3A**). This suggests that there are other features that impact the rates, e.g., the shape of the membrane distortion, groove dynamics, and residues lining the groove. Another feature we have not explored are mixed membranes and membranes of shorter or longer chain length which we expect would alter lipid scrambling rates. For example, TMEM16K resides in the endoplasmic reticulum (ER) membrane which is thinner than the plasma membrane (12, 33, 80), and TMEM16K scrambling rates increase tenfold in thinner membranes (12).

Experimental scrambling assays performed by different groups have reported basal level scramblase activity in the absence of Ca^2+^ for fungal and mammalian dual-function scramblases (11–13, 19, 40, 43, 57). It is unknown where closed-groove scrambling takes place on the protein (62) and simulations have never reported such events despite Li and co-workers reporting scrambling events for simulations initiated from closed TMEM16A, TMEM16K, and TMEM16F (79), which may have also been sampled in these trajectories. In aggregate, we observed 60 scrambling events that do not follow the credit-card model and occur “out-of-the-groove” (**Table 1**). Nearly all these events (56/60) happen at the dimer interface between TM3 and TM10 of the opposite subunit, here on referred to as the dimer cleft. Curiously, we do not observe scrambling at this location for any of the fungal structures. Although mammalian TMEM16s have a ∼4-5 Å wider gap on average at the lower leaflet dimer cleft entrance than the open fungal TMEM16s, we do not always observe scrambling at such distances and sometimes do not observe any scrambling when the cleft is at its widest (**Appendix 1-Figure 7**). For all structures we see lipids from both leaflets intercalate between TM3 and TM10 (**Figure 4-figure supplement 6**), which is consistent with lipid densities in cryo-EM nanodiscs images of fungal TMEM16s (59, 62) and TMEM16F (66). Based on our simulations, this interface may be a source for Ca^2+^-independent scrambling.

It is unclear whether the out-of-the-groove events we have observed reflect the same closed-groove scrambling activity seen in experimental assays (40, 43, 59, 62). Also, it is possible that we have missed slow or rare out-of-the-groove events due to limited sampling. One way to assess these points is to ask whether the relative scrambling rates observed in +/- Ca^2+^ are similar to the relative rates from our simulation with open/closed hydrophilic grooves. Feng *et al.* reported a 7-18 fold increase in scrambling rate by nhTMEM16 in the presence of Ca^2+^ compared to Ca^2+^-free conditions (62). Based on our open groove count of 220, we would expect 12-30 events for the closed groove states, but we observed no events. However, Watanabe and colleagues reported a 6-7 fold increase in scrambling rate by TMEM16F in the presence of Ca^2+^ compared to Ca^2+^-free conditions (57), which is consistent with the 7-9 fold increase revealed in our simulations between closed Ca^2+^-free TMEM16F structures (PBD IDs 6P47 and 6QPB) and the WT open Ca^2+^-bound TMEM16F (6QP6*). It is possible that out-of-the-groove scrambling is highly dependent on the membrane composition, as discussed earlier, and the scrambling ratios we observe in DOPC may be different than the experimental rates determined in different lipids. This cannot be addressed without additional studies. That said, we are encouraged by the high-level correspondence in TMEM16F – we observe much higher scrambling rates through the open grooves and much smaller flipping rates elsewhere on the protein or with closed groove structures, suggesting that our simulations may be revealing aspects of Ca^2+^-independent scrambling in mammalian family members.

With regard to predicting absolute rates, our simulations correctly distinguish scrambling competent structures from non-competent scramblers, but direct comparison of our rates with experimental values (that tend to be 2-3 orders of magnitude slower) should be interpreted qualitatively. For example, single-molecule analysis yielded a scrambling rate of 0.04 events per μs for TMEM16F (57), whereas we find 3 and 11.3 events per μs for our scrambling-competent TMEM16F structures 6QP6* and PDB ID 8B8J, respectively. Malvezzi and co-workers estimated a similar scrambling rate of 0.02 events per μs for afTMEM16 using a liposome-based assay while we find 10.7 events per μs (43). We will highlight three potential explanations for such discrepancy. First, it is well established that the Martini model increases diffusion dynamics by a factor ∼4 due to the lower friction between CG beads and reduced configurational entropy compared to more chemically detailed representations (81). Second, the energy barrier for a PC headgroup to traverse the DOPC bilayer in absence of protein is reduced in Martini 3 compared to Martini 2 and AAMD (82). It is not trivial to predict how this reduction affects protein-mediated lipid scrambling, but it is likely to increase observed flipping rates compared to the more realistic AAMD. Third, as shown in **Fig. 3*B***, the Martini 3 elastic network used to restrain the protein backbone in our simulations allows a small degree of flexibility during simulations, which may increase scrambling. For instance, the groove of the open nhTMEM16 structure 4WIS enlarges by ∼3 Å during our Martini 3 simulations compared to the starting experimental structure and our Martini 2 simulations, and this dilation correlates with greater scrambling (**Appendix 1-Figure 8A-B**). We also analyzed previously published CHARMM36 AAMD trajectories starting from the same structure (36) and observed that while these simulations do show some degree of dilation, as we observe with Martini 3, they generally stay closer to the experimental structure (**Appendix 1-Figure 8B**). In addition to the open nhTMEM16 structure, we observed similar subtle movements in the TM4 helix for open Ca^2+^-bound structures of afTMEM16, TMEM16K, TMEM16F, and TMEM16A that appear to enlarge the TM4/TM6 outer vestibule (**Figure 3-figure supplement 6A**). Others have reported that AAMD simulations sample spontaneous dilation of the groove/pore to confer either scramblase activity for WT (52, 55) and mutant (58) TMEM16F or ion channel activity for TMEM16A structures (47, 50, 83). These movements away from the experimentally solved structures may be due to the inaccuracy of our atomistic and CG force fields or differences in the model and experimental membrane/detergent environments, but more work is needed to assess whether these dilations reflect physiologically relevant conformational states. CG simulations of closed-groove structures lack such dilations, because the backbones of TM4 and TM6 are in close enough proximity (< 10 Å) to be connected by the elastic network that the Martini model requires to maintain proper secondary and tertiary structure (e.g., PDB ID 6QM4, see **Appendix 1-Figure 8C-D**). The recent GōMartini 3 model replaces the harmonic bonds of the elastic network with Lennard-Jones potentials that vanish as residues separate potentially making this an excellent model for sampling groove opening and closing (84) (**Appendix 1-Figure 8E**).

Finally, we end by discussing the observed in- and out-of-the-groove scrambling for the putative ion conducting states of TMEM16A. The low number of recorded events (11 for the highest and 1 for the lowest) may be consistent with the lack of experimentally measured scramblase activity (54, 70), for the reasons discussed in the last paragraph. Consistent with our low computational rate, we also computed an energy barrier for lipid movement through the TMEM16A groove 5.5-fold higher than the scramblase barriers (**Appendix 1-Figure 3B**). In simulations of our three predicted ion conductive states of TMEM16A (7ZK3*^6,8,10^) lipid headgroups insert into the lower and upper vestibule of the pore. Compared to the inhibitor-bound structure (PDB ID 7ZK3), the outer vestibule of these conductive states is notably more dilated. We observe 4 Cl^-^ permeation events by 7ZK3*^8^ through the partially lipid-lined groove. Surprisingly, our simulation of the predicted TMEM16A conductive state from Jia & Chen did at times feature a fully-lipid lined groove, similar to the proteolipidic pore found in dual-function members (48) (**Fig. 1*D***, **Figure 3-video 1**); however, we did not observe any ion permeation events from this configuration, which may be a consequence of the configuration not being physiologically relevant, the Martini 3 force field not being ideal for Cl^-^/lipid/protein interactions, or something else. It is intriguing that while TMEM16A has lost experimentally discernable scrambling activity, it still deforms and thins the membrane (**Fig. 3A**). Coupled with our observation that groove widening allows lipids to enter, we wonder if it retains thinning capabilities to facilitate partial lipid insertion to promote Cl^-^ permeation. This hypothesis has been stated before (85), and structural evidence for this proteolipidic ion channel pore has recently been reported for the OSCA1.2 mechanosensitive ion channel, which adopts the TMEM16 fold, yet it does not scramble lipids (51, 86).

## Materials and Methods

### Coarse-grained system preparation and simulation details

For each simulated structure, missing loops with less than 16 residues were modeled using the loop building and refinement procedures MODELLER (version 10.2 (87)). Further details on which loops were included are in **Appendix 1 – Table 1**. For each stretch of *N* missing residues 10x*N* models were generated. We then manually assessed the 10 lowest DOPE scoring predictions and selected the best model based on visual inspection. Models were inserted symmetrically into the original experimental dimer structure except for PDB IDs 8BC0, 8TAG, and 5OC9 which were published as asymmetric structures.

Setup of the CG simulation systems was automated in a python wrapper script adapted from MemProtMD (35). After preparing the atomistic structure using pdb2pqr (88), the script predicted protein orientation with respect to a membrane with memembed (89). Then, martinize2 (90) was employed to build a Martini 3 CG protein model. Secondary structure elements were predicted by DSSP (91) and their inter- and intra-orientations within a 5-10 Å distance were constrained by an elastic network with a 500 kJ mol^-1^ nm^-2^ force constant (unless specified otherwise). CG Ca^2+^ ions (bead type “SD” in Martini 3) were inserted at their respective positions based on the original protein structure and connected to coordinating (<= 6 Å) Asp and/or Glu side chains by a harmonic bond with a 100 kJ mol^-1^ nm^-2^ force constant. A DOPC membrane was built around the CG protein structure using *insane* (92) in a solvated box of 220×220×180 Å^3^, with 150 mM NaCl. Systems were charge-neutralized by adding Cl^-^ or Na^+^ ions. For each system, energy minimization and a 2 ns NPT equilibration were performed. All systems were simulated for 10 μs in the production phase and the first microsecond was excluded from all analyses for equilibration.

All CG molecular dynamics simulations were performed with Gromacs (version 2020.6 (93)) and the Martini 3 force field (version 3.0.0 (73)). A 20 fs time step was used. Reaction-field electrostatics and Van der Waals potentials were cut-off at 1.1 nm (94). As recommended by Kim *et al.* (95), the neighbor list was updated every 20 steps using the Verlet scheme with a 1.35 nm cut-off distance. Temperature was kept at 310 K using the velocity rescaling (96) thermostat (τ_T_=1 ps). The pressure of the system was semi-isotropically coupled to a 1 bar reference pressure by the Parrinello-Rahman (97) barostat (τ_P_=12 ps, compressibility=3×10^-4^).

### Lipid headgroup and water density calculations

First, each protein subunit was individually aligned in x, y, and z to their starting coordinates. Atomistic simulations were filtered for trajectory frames with T333-Y439 Cα distance >15 Å giving a total of ∼2085 ns of aggregate simulation time. Then the positions of all PC headgroup beads were tracked overtime and binned in a 100×100×150 Å grid with 0.5 Å spacing centered on two residues near the membrane midplane on TM4 and TM6 using a custom script that includes MDAnalysis methods (98, 99). Density for water beads was calculated in the same way. Density in each cell was then averaged from each chain and for atomistic simulations averaged from all 8 independent simulations.

### Scrambling analysis

Lipid scrambling was analyzed as described by Li *et al.* (71). For every simulation frame (1 ns^-1^ sampling rate), the angle between each individual DOPC lipid and the z-axis was calculated using the average of the vectors between the choline (NC3) bead and the two last tail beads (C4A and C4B), see **Figure 2-figure supplement 1A**. We applied a 100 ns running average to denoise the angle traces. Lipids that reside in the upper leaflet are characterized by a 150° angle, and lipids in the lower leaflet have a 30° angle. Scrambling events were counted when a lipid from the upper leaflet passed the lower threshold at 35° or, *vice versa*, when a lipid from the lower leaflet passed the upper threshold at 145° (see **Figure 2-figure supplement 1B**). These settings are more stringent than the thresholds used by Li *et al.* (55° and 125°, respectively) to prevent falsely counted partial transitions (70). A 1 μs block averaging was applied to obtain averages and standard deviations for the scrambling rates.

### Groove dilation analysis

The residues chosen for measuring the minimum distance between TM4 and TM6 were located within ∼6 Å in z (1-2 α-helix turns) of the path node with the minimum net flux of water (see Appendix 1-Methods). The residues used for each homolog were as follows: 327-339 and 430-452 for nhTMEM16, 319-331 and 426-438 for afTMEM16, 365-377 and 434-446 for TMEM16K, 512-424 and 613-625 for TMEM16F, and 541-553 and 635-647 for TMEM16A. Distances were calculated using a custom script that includes MDAnalysis methods (98, 99).

### Quantification of membrane deformations

First, using Gromacs (gmx trjconv), MD trajectories were aligned in the xy-plane such that the longest principal axis defined by the initial positions of TM7 and TM8 aligned to the global y axis. Average membrane surfaces were calculated from the aligned MD trajectories as outlined previously (36) using a custom python script based on MDAnalysis (98) and SciPy (100). The positions of each lipid’s glycerol beads (GL1 and GL2) were linearly interpolated to a rectilinear grid with 1 Å spacing. Averaging over all time frames (again, discarding the first 1 μs for equilibration) yielded a representative upper and lower leaflet surface. Grid points with a lipid occupancy below 2% were discarded. Clusters of grid points that were disconnected from the bulk membrane surface were discarded. The minimal membrane thickness was calculated as the minimal distance between any two points on the opposing ensemble-averaged surfaces (e.g., **Fig. 3C**). Crucially, in the case of lipid scrambling simulations like the ones described here, lipids were assigned to the upper/lower leaflet separately for every time frame.

### Protein-lipid contact and dwell time analysis

Using the full 10 μs simulation where each protein subunit was individually aligned in x, y, and z, we analyzed protein-lipid interactions by measuring distances between the protein’s outermost sidechain bead (except for glycine, which only has backbone bead) and the lipid’s choline (NC3) or phosphate (PO4) bead for every nanosecond using custom scripts with Scipy methods (100). Contacts were defined as distances below 7 Å. Contact frequency was calculated as the fraction of simulation frames where a contact occurred, averaged over two monomers. Dwell time was measured as the duration of consecutive contacts, allowing breaks up to 6 ns to account for transient fluctuations of lipid configuration. For each residue, we selected either the choline or phosphate bead based on which yielded the higher average dwell time. To visualize the result, we used averaged dwell time of the top 50% longest dwelling events at each residue to generate a color-coded representation of the protein structure (**Fig. 4C-D**; **Figure 4-figure supplement 3**).

### Simulation and data visualization

Each simulation video and all simulation snapshots with lipid headgroup coordinate densities and traces, average membrane surfaces, and protein colored by lipid contact/dwell time were rendered using VMD (101). Images of TMEM16A atomistic starting structures were rendered using ChimeraX (102). All plots were generated using the Matplotlib graphics package (103).

## Data Availability Statement

All code used to generate main figures and analyze MD trajectories as well as original MD trajectories will be made available upon request to the corresponding author.

## Acknowledgements

Neville Bethel performed the all-atom simulations of nhTMEM16 reported in ref. (36). Paola Bisignano performed the loop modeling for many of the TMEM16F structures. George Khelashvili generously shared the simulated open-groove structure of TMEM16F from ref. (55). We thank Wenlei Ye for his helpful comments on our manuscript. We thank Andrew Natale and Yessica Gomez who helped develop the membrane deformation analysis scripts.

## Funding information

This work was supported by funding from National Institutes of Health (NIH) Grant R01 GM137109 (MG), National Science Foundation (NSF) award MCB-2217662 (MG), UCSF Discover Fellows Program (CAS), and National Science Foundation Graduate Research Fellowship Program Grant No. 2038436 (CAS).

## Figure Supplements

**Figure 1-figure supplement 1.**
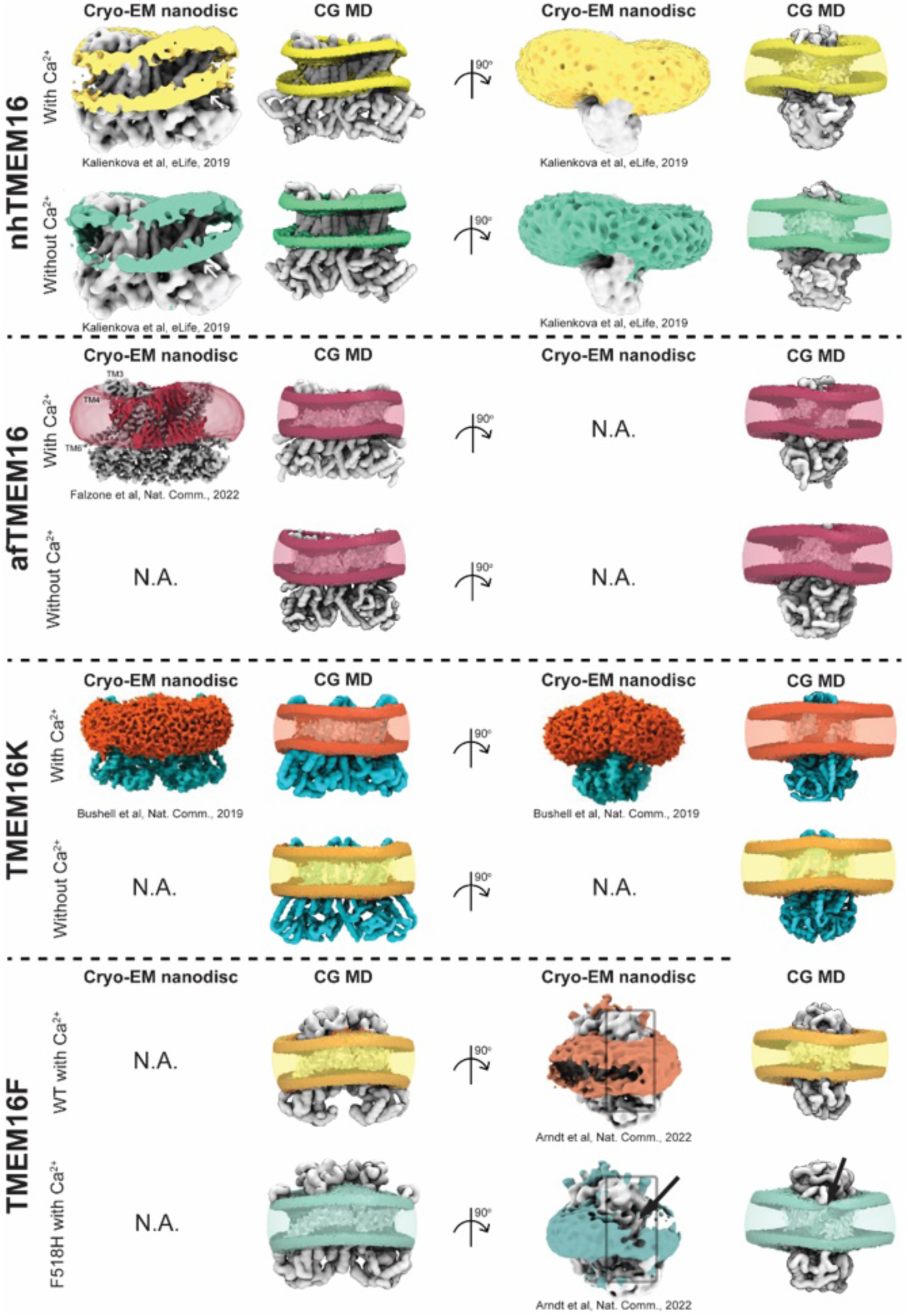
Comparison between membrane deformations in cryo-EM nanodiscs and CG MD. Front and side views of cryo-EM maps (left) and ensemble-averaged conformations from CG MD simulations (right). nhTMEM16: cryo-EM images (PDB IDs 6QM9 and 6QM4) were adapted from © 2019, Kalienkova et al, published by eLife (61). The CG MD structures are PDB IDs 4WIS (with Ca^2+^) and 6QM4 (without Ca^2+^). afTMEM16: cryo-EM image (PDB ID 7RXG) was adapted from © 2022, Falzone et al, published by Springer Nature (59). The CG MD structures are PDB IDs 7RXG (with Ca^2+^) and 7RXB (without Ca^2+^). TMEM16K: cryo-EM images (PDB ID 5OC9) were adapted from © 2019, Bushell et al, published by Springer Nature (12). The CG MD structures are PDB IDs 5OC9 (with Ca^2+^) and 6R7X (without Ca^2+^). TMEM16F: cryo-EM images (PDB IDs 6QPC and 8B8J) were adapted from © 2022, Arndt et al, published by Springer Nature (60). The CG MD structures are also PDB IDs 6QPC (WT with Ca^2+^) and 8B8J (F518H with Ca^2+^). Black arrows indicate a dip in lipid density near the groove entrance. CG MD images show the average glycerol and headgroup densities in solid and the tail densities in transparent coloring that matches the colors used in the original cryo-EM images.

**Figure 1-figure supplement 2.**
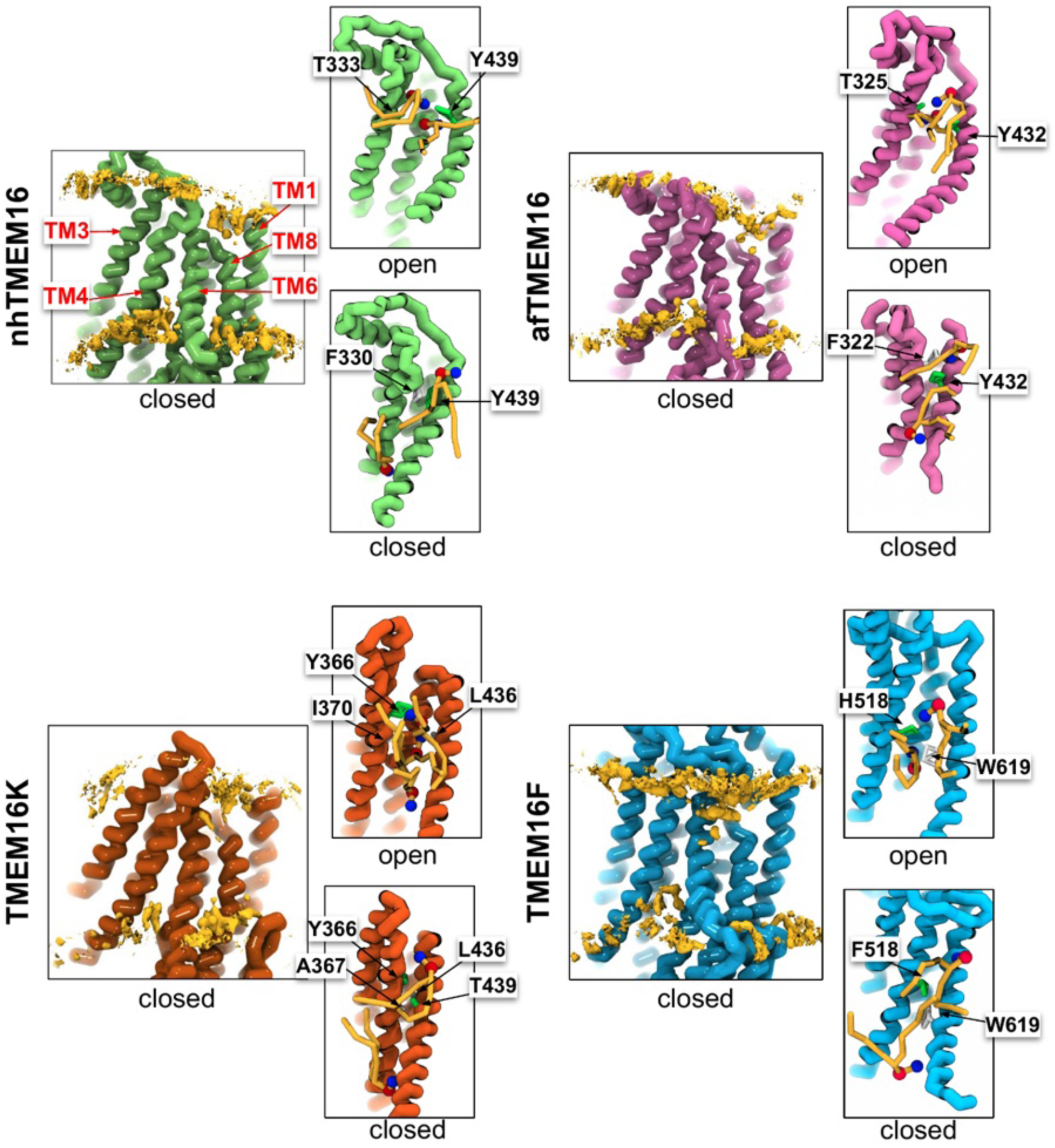
CG simulations of multiple TMEM16 structures with closed grooves lack lipid density in the TM4-TM6 pathway. Snapshots from CG MD simulations of nhTMEM16 (PDB IDs 6QM4 (closed) and 4WIS (open), green), afTMEM16 (PDB IDs 7RXB (closed) and 7RXG (open), violet), TMEM16K (PDB IDs 6R7X (closed) and 5OC9 (open), gold), and TMEM16F (PDB IDs 6QPB (closed) and 6QP6* (open), blue) with phosphatidylcholine (PC) lipid headgroup density (yellow) and nearby lipids (yellow). Residues forming the closest distance between TM4 and TM6 (colored by residue type: basic (red), acidic (blue), and polar (green)) and lipids near the groove also shown. Each density is averaged over both chains.

**Figure 1-table supplement 1.**
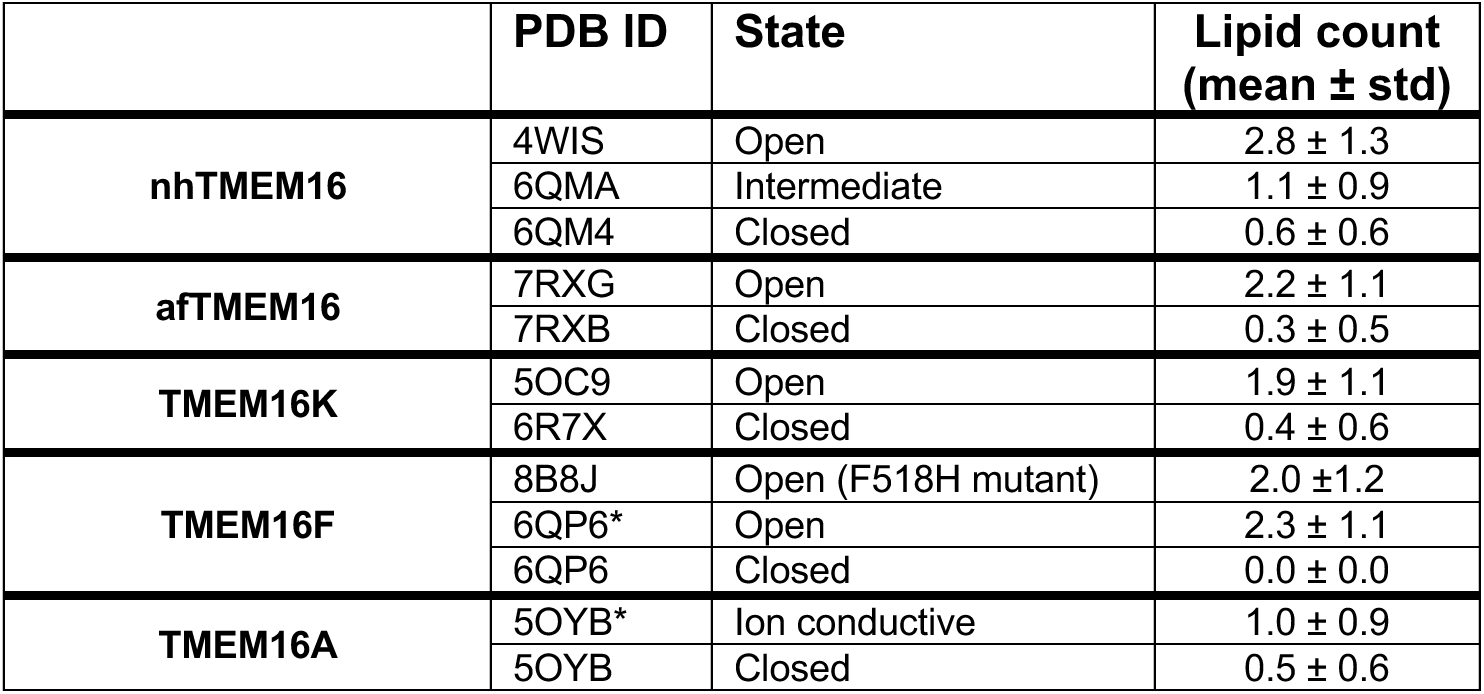
Lipid counts in the TM4/TM6 groove. At every simulation frame, the number of lipid headgroups in the groove was counted, based on their z-position.

**Figure 2-figure supplement 1.**
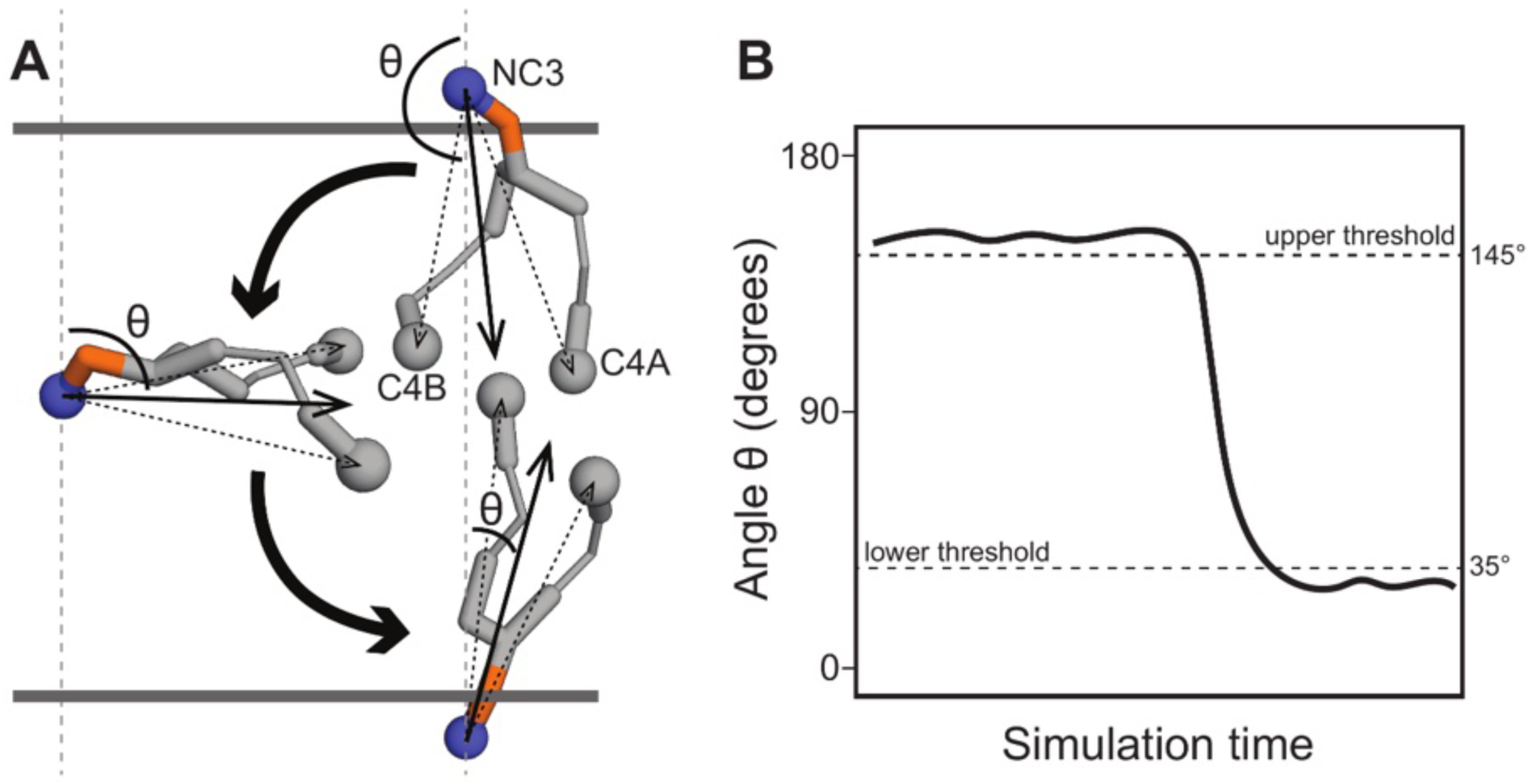
Measuring lipid angles to detect scrambling events. **(A)** For every time frame, for every lipid in the system, we defined a vector between the choline bead (NC3) and the two tail beads (C4A, C4B; dashed arrows) and calculated the angle θ between the average of those two vectors (solid arrow) with the z axis. **(B)** A schematic representation of a typical time trace for a lipid that scrambles from the upper membrane leaflet ( θ ≈ 150°) to the lower membrane leaflet (θ ≈ 30°). A scrambling event is only counted when θ passes the threshold at the opposite leaflet with respect to its original location (35° for the lower, 145° for the upper).

**Figure 2-figure supplement 2.**
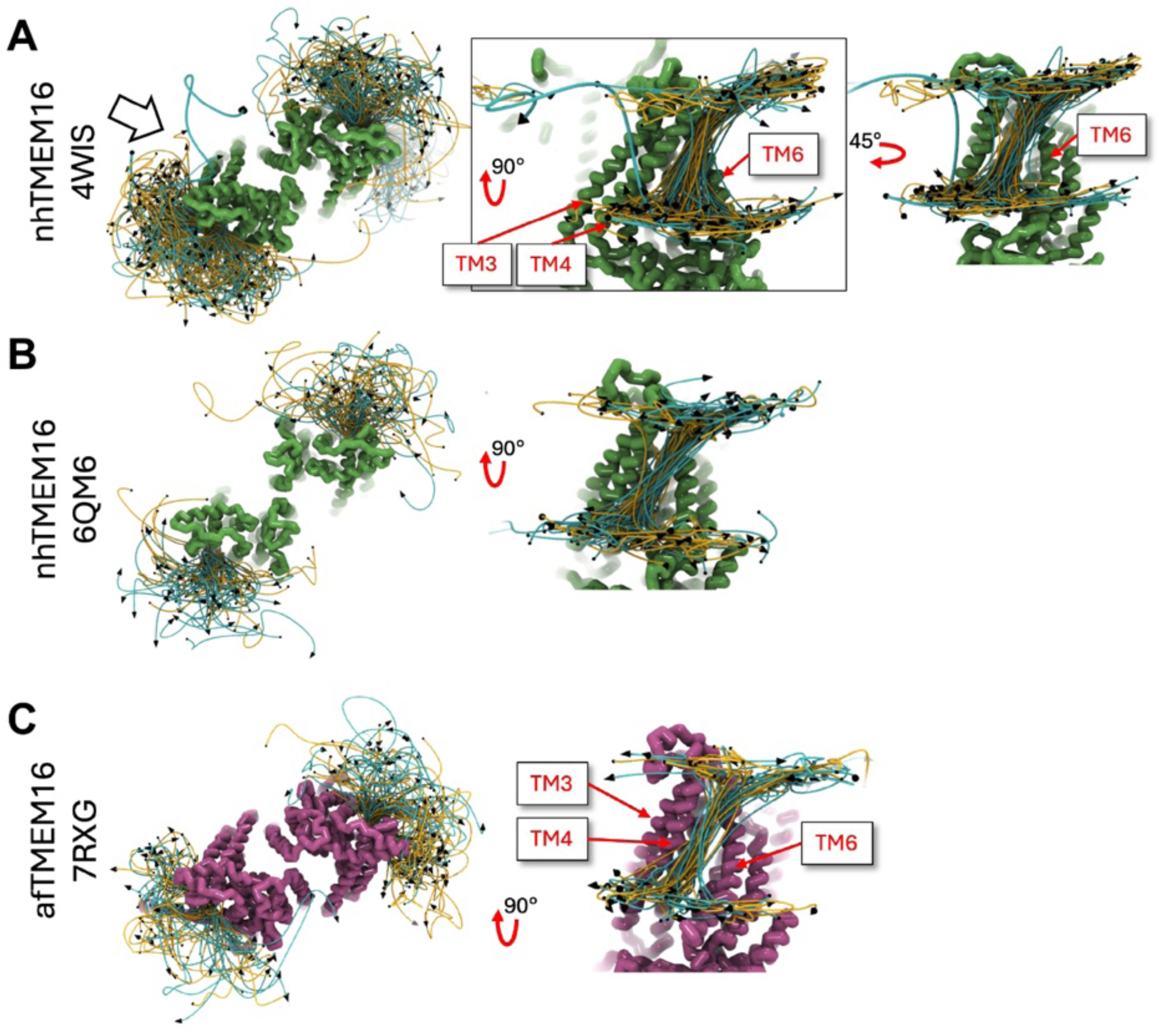
Position traces of scrambling lipids in fungal TMEM16 simulations. **(A)** Lipid traces for Ca^2+^-bound open nhTMEM16 (PDB ID 4WIS). **(B)** Lipid traces for Ca^2+^-bound open nhTMEM16 (PDB ID 6QM6). **(C)** Lipid traces for Ca^2+^ -bound open afTMEM16 (PDB ID 7RXG). Lipid traces are generated by fitting raw lipid headgroup center of mass positions to a smooth spline curve.

**Figure 2-figure supplement 3.**
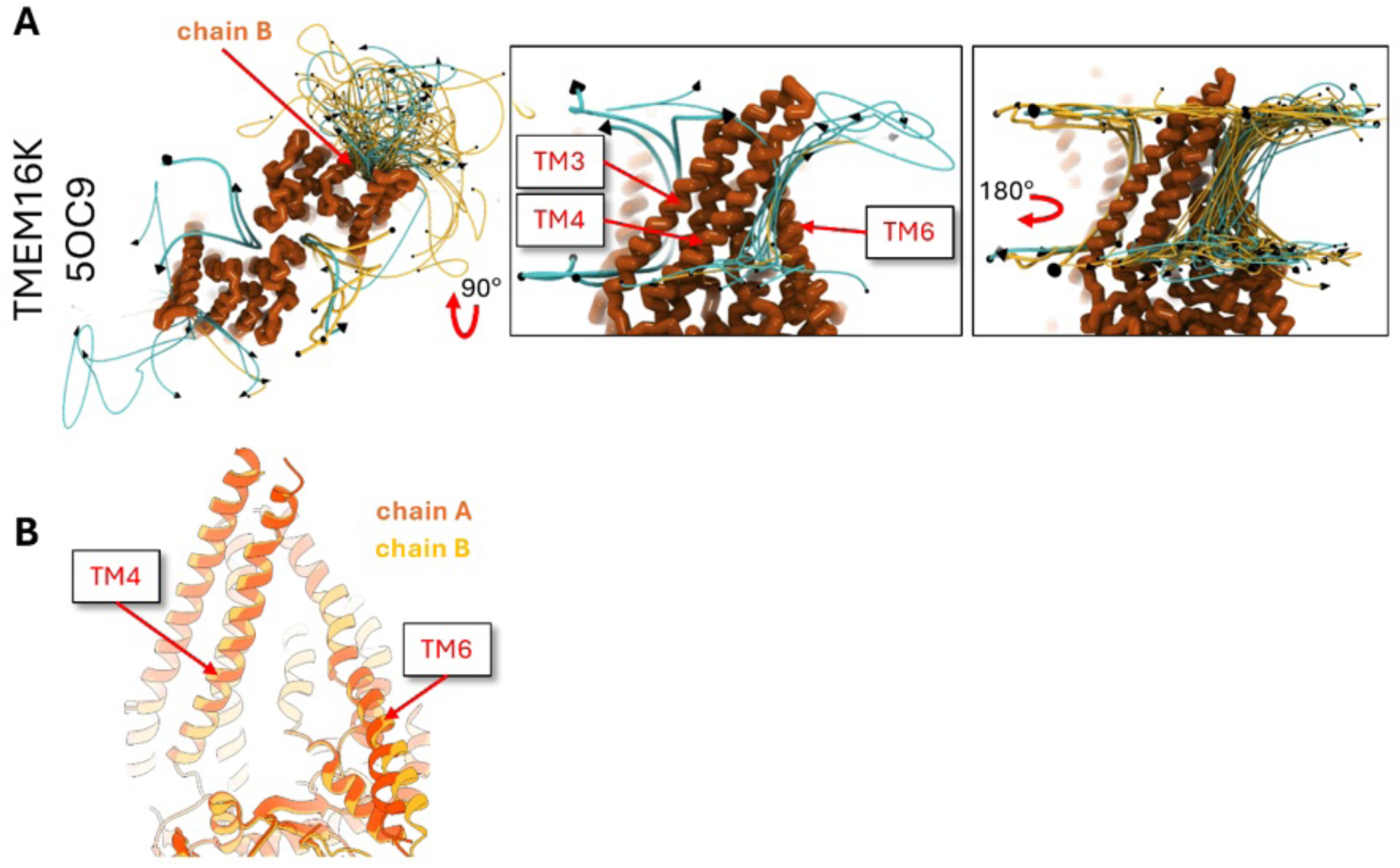
Position traces of scrambling lipids in the open TMEM16K simulation. **(A)** Lipid traces for Ca^2+^-bound open TMEM16K (PDB ID 5OC9). **(B)** Cartoon representation of aligned subunits of Ca^2+^-bound TMEM16K (PDB ID 5OC9). Lipid traces are generated by fitting raw lipid headgroup center of mass positions to a smooth spline curve.

**Figure 2-figure supplement 4.**
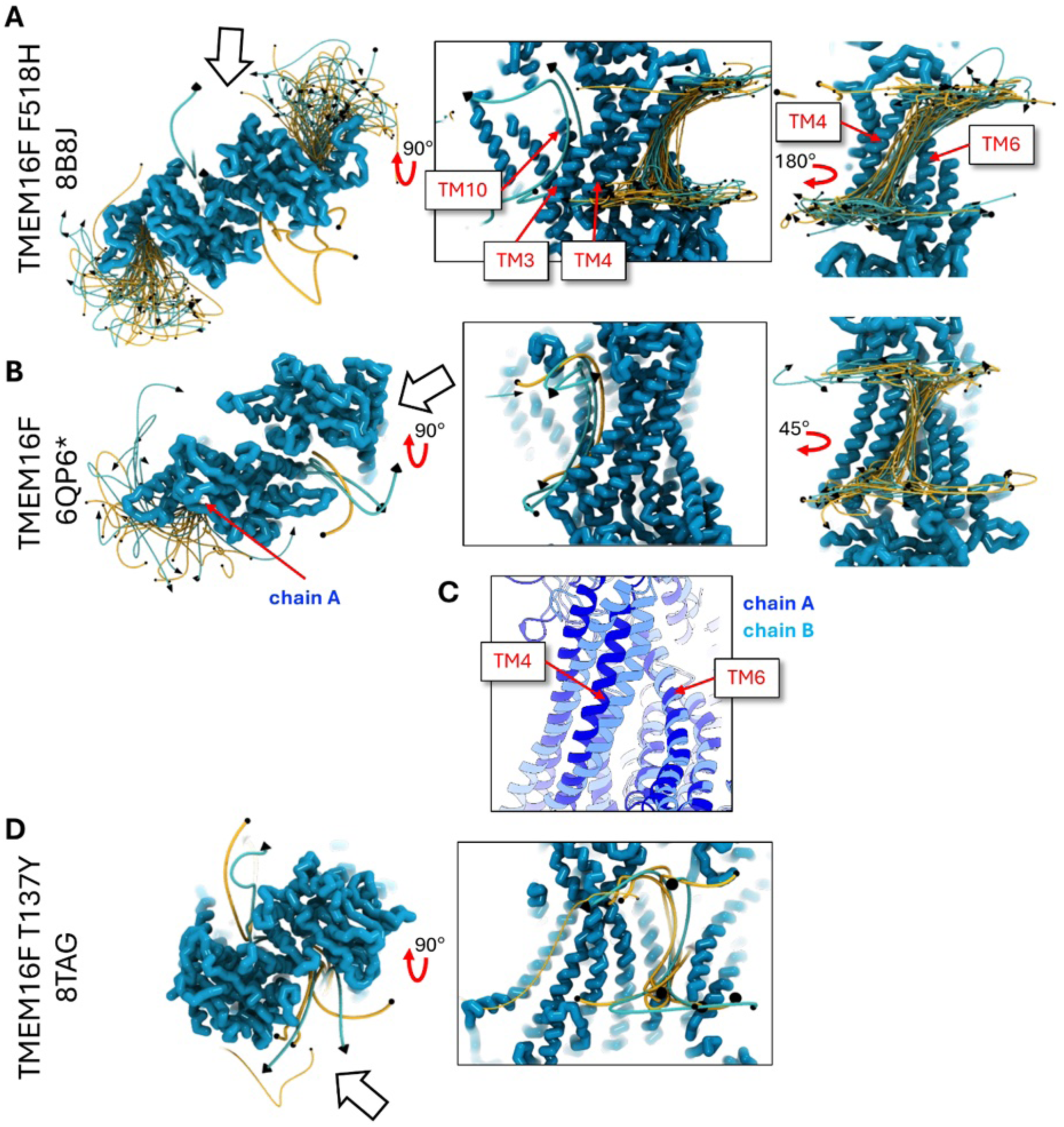
Position traces of scrambling lipids in TMEM16F simulations. **(A)** Lipid traces for Ca^2+^ -bound open TMEM16F F518H mutant (PDB ID 8B8J). **(B)** Lipid traces for Ca^2+^ -bound simulated open TMEM16F (6QP6*) **(C)** Cartoon representation of aligned subunits of Ca^2+^-bound simulated open TMEM16F (6QP6*). **(D)** Lipid traces for Ca^2+^ -bound simulated closed TMEM16F (PDB ID 8TAG). Lipid traces are generated by fitting raw lipid headgroup center of mass positions to a smooth spline curve.

**Figure 2-figure supplement 5.**
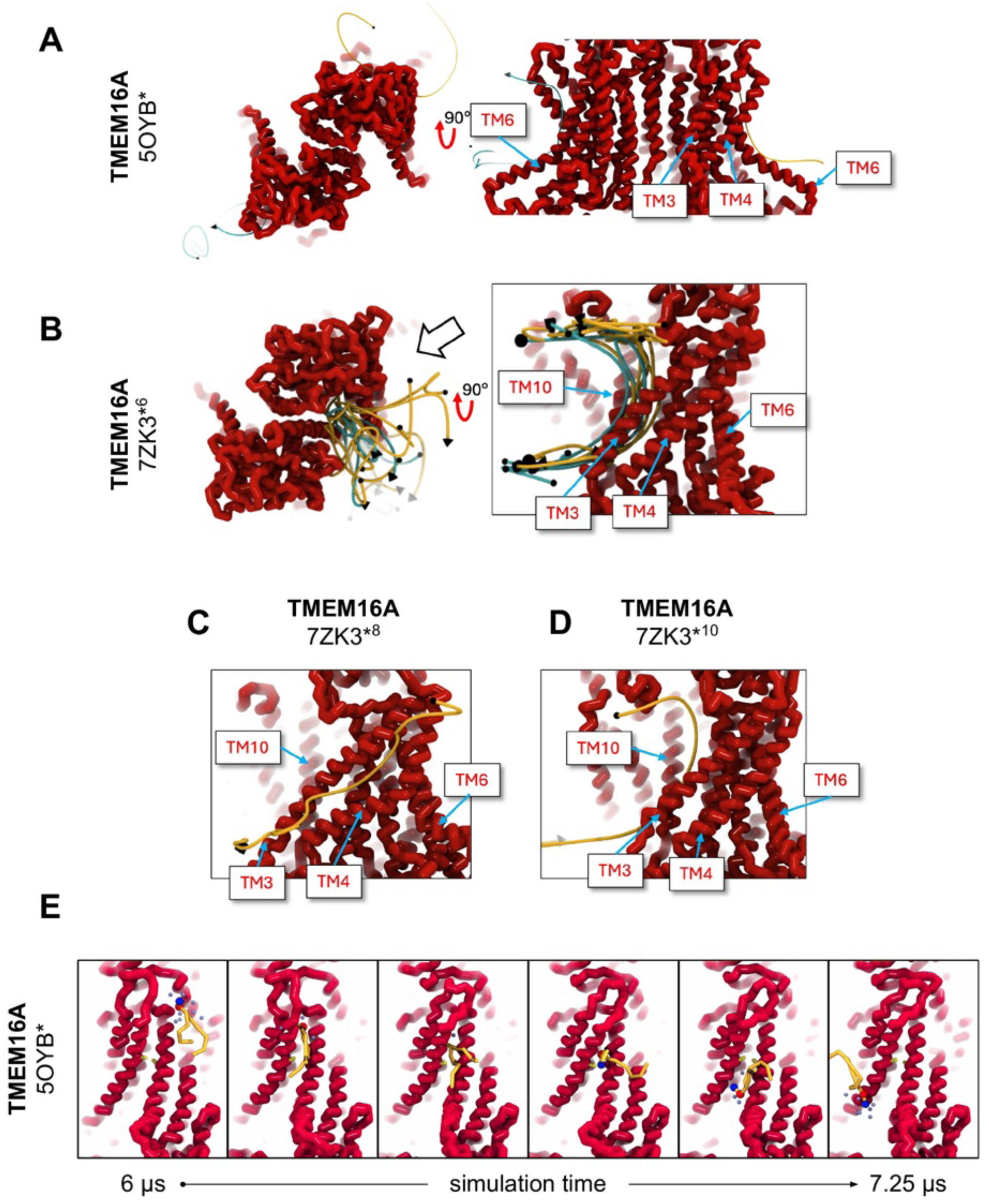
Position traces of scrambling lipids in TMEM16A simulations. **(A)** Lipid traces for Ca^2+^ -bound simulated conductive TMEM16A (5OYB*). **(B)** Lipid traces for Ca^2+^ -bound simulated open TMEM16A (7ZK3*^6^). **(C)** Lipid traces for Ca^2+^ -bound simulated open TMEM16A (7ZK3*^10^). **(D)** Lipid traces for Ca^2+^ -bound simulated open TMEM16A (7ZK3*^8^). **(E)** Snapshots from simulation of 5OYB* during lipid scrambling event (trace in **A**). Lipid traces are generated by fitting raw lipid headgroup center of mass positions to a smooth spline curve.

**Figure 3-figure supplement 1.**
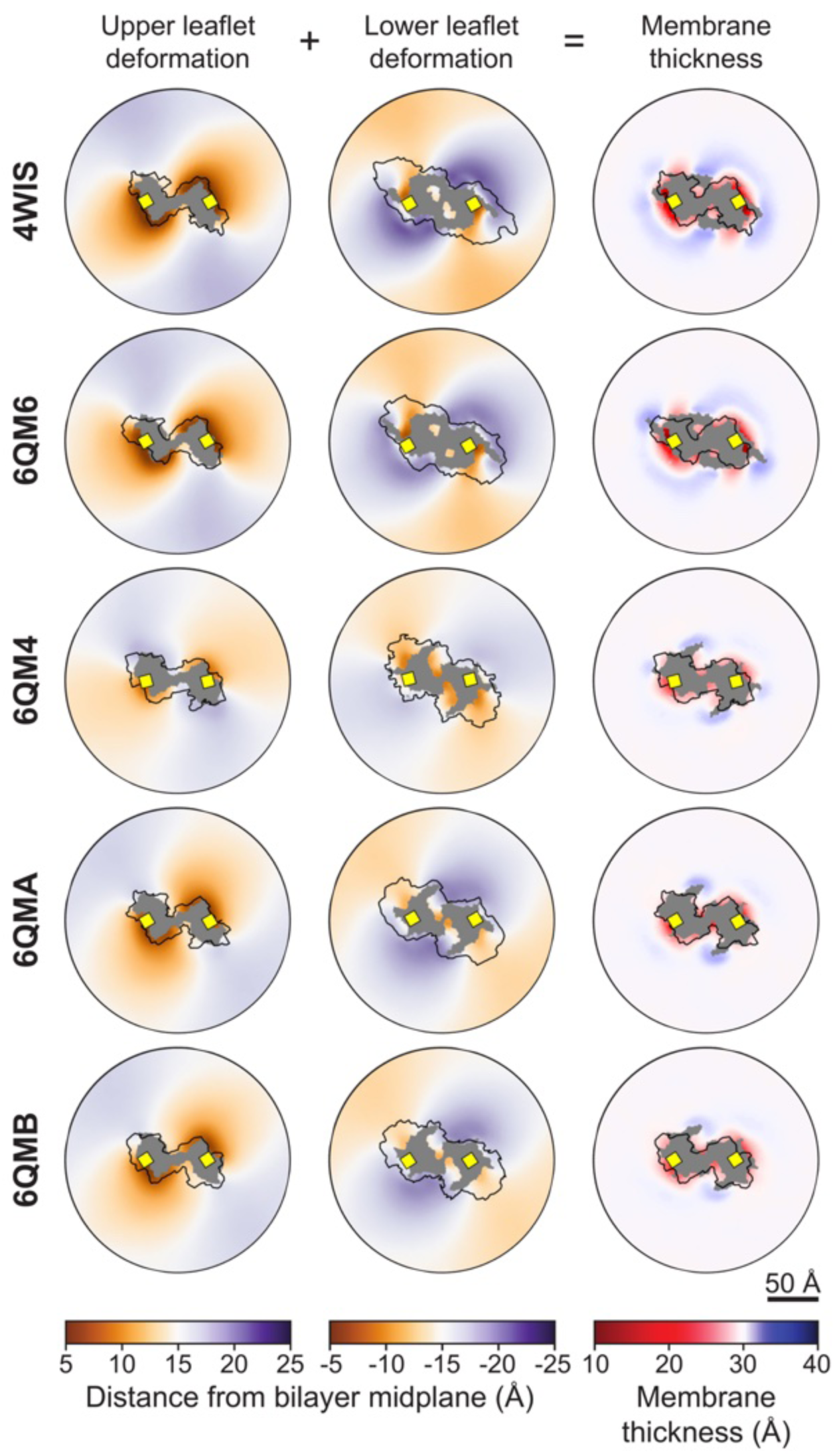
Membrane deformations for simulated nhTMEM16 structures. Left column: xy-map of the distance along the z-axis from the bilayer midplane to the ensemble averaged positions of the glycerol linker (GL1 and GL2 beads). Middle column: xy-map of the distance along the z-axis from the bilayer midplane to the ensemble averaged positions of the glycerol linker (GL1 and GL2 beads). Right column: the sum of the upper and lower leaflet deformations, representing the bilayer thickness along z. In all plots, grey areas indicate grid points with lipid occupancy <2%. The black outline is the projected surface of the upper (z>0) or lower (z<0) portion of the protein dimer.

**Figure 3-figure supplement 2.**
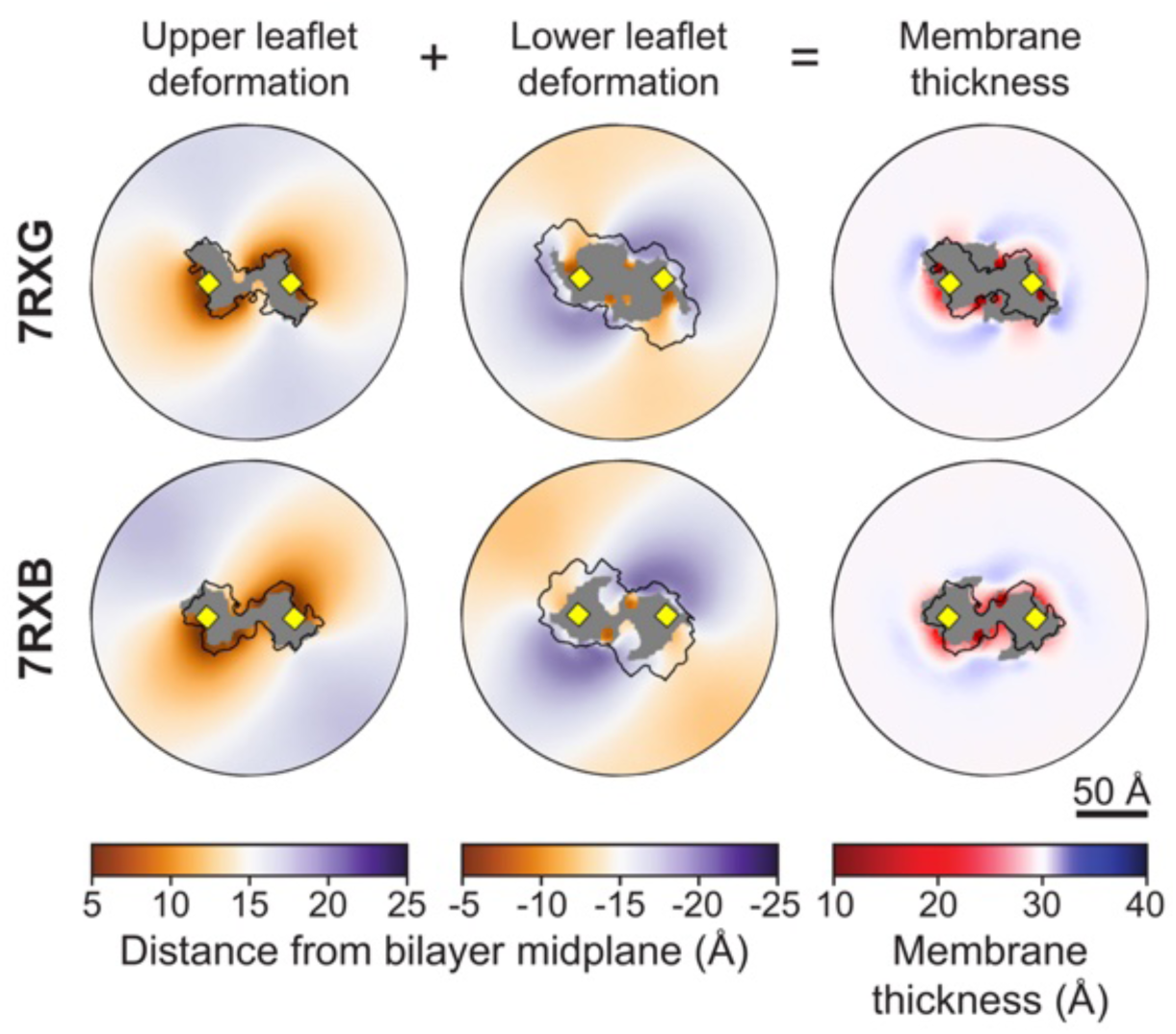
Membrane deformations for simulated afTMEM16 structures. Left column: xy-map of the distance along the z-axis from the bilayer midplane to the ensemble averaged positions of the glycerol linker (GL1 and GL2 beads). Middle column: xy-map of the distance along the z-axis from the bilayer midplane to the ensemble averaged positions of the glycerol linker (GL1 and GL2 beads). Right column: the sum of the upper and lower leaflet deformations, representing the bilayer thickness along z. In all plots, grey areas indicate grid points with lipid occupancy <2%. The black outline is the projected surface of the upper (z>0) or lower (z<0) portion of the protein dimer.

**Figure 3-figure supplement 3.**
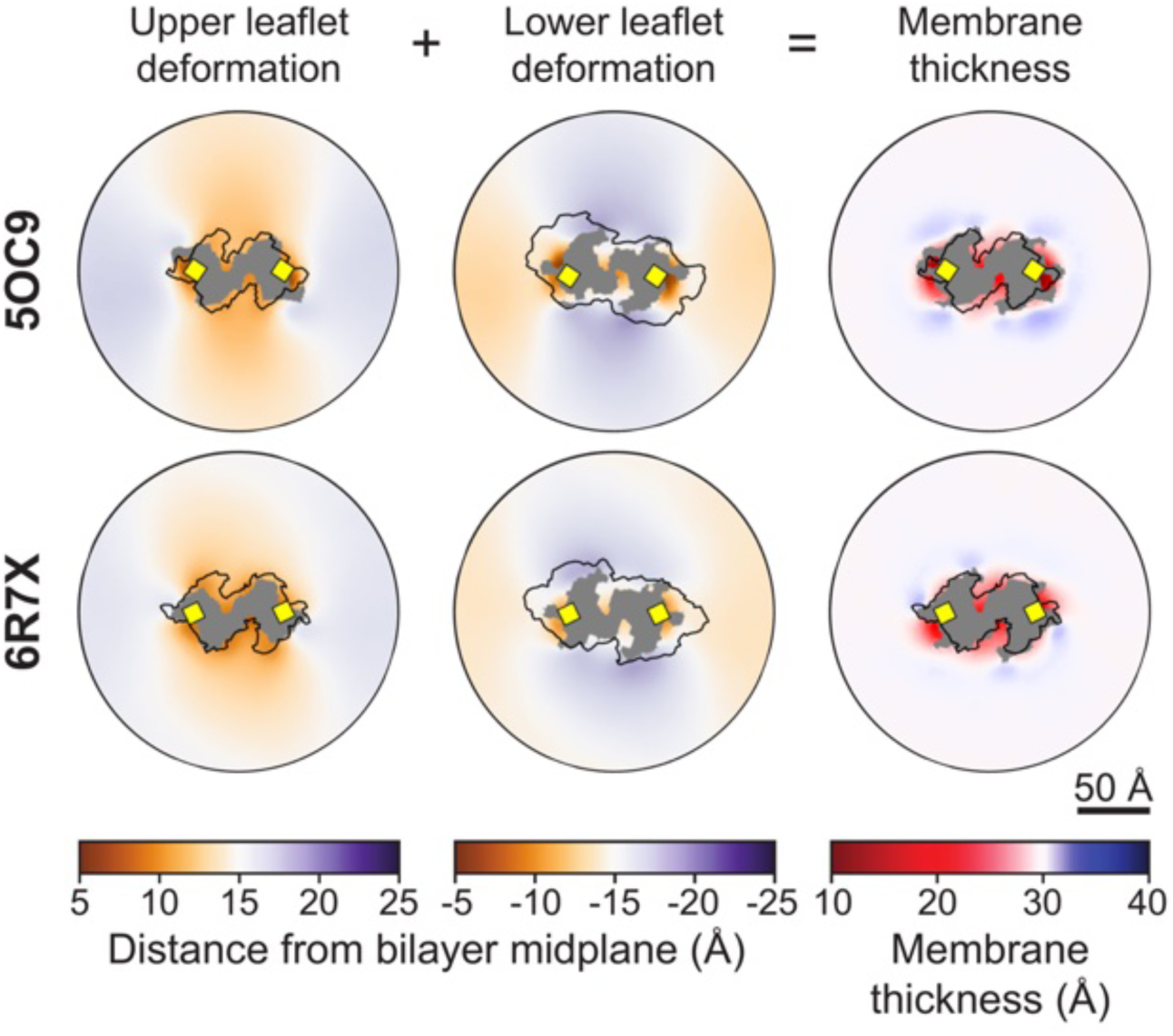
Membrane deformations for simulated TMEM16K structures. Left column: xy-map of the distance along the z-axis from the bilayer midplane to the ensemble averaged positions of the glycerol linker (GL1 and GL2 beads). Middle column: xy-map of the distance along the z-axis from the bilayer midplane to the ensemble averaged positions of the glycerol linker (GL1 and GL2 beads). Right column: the sum of the upper and lower leaflet deformations, representing the bilayer thickness along z. In all plots, grey areas indicate grid points with lipid occupancy <2%. The black outline is the projected surface of the upper (z>0) or lower (z<0) portion of the protein dimer.

**Figure 3-figure supplement 4.**
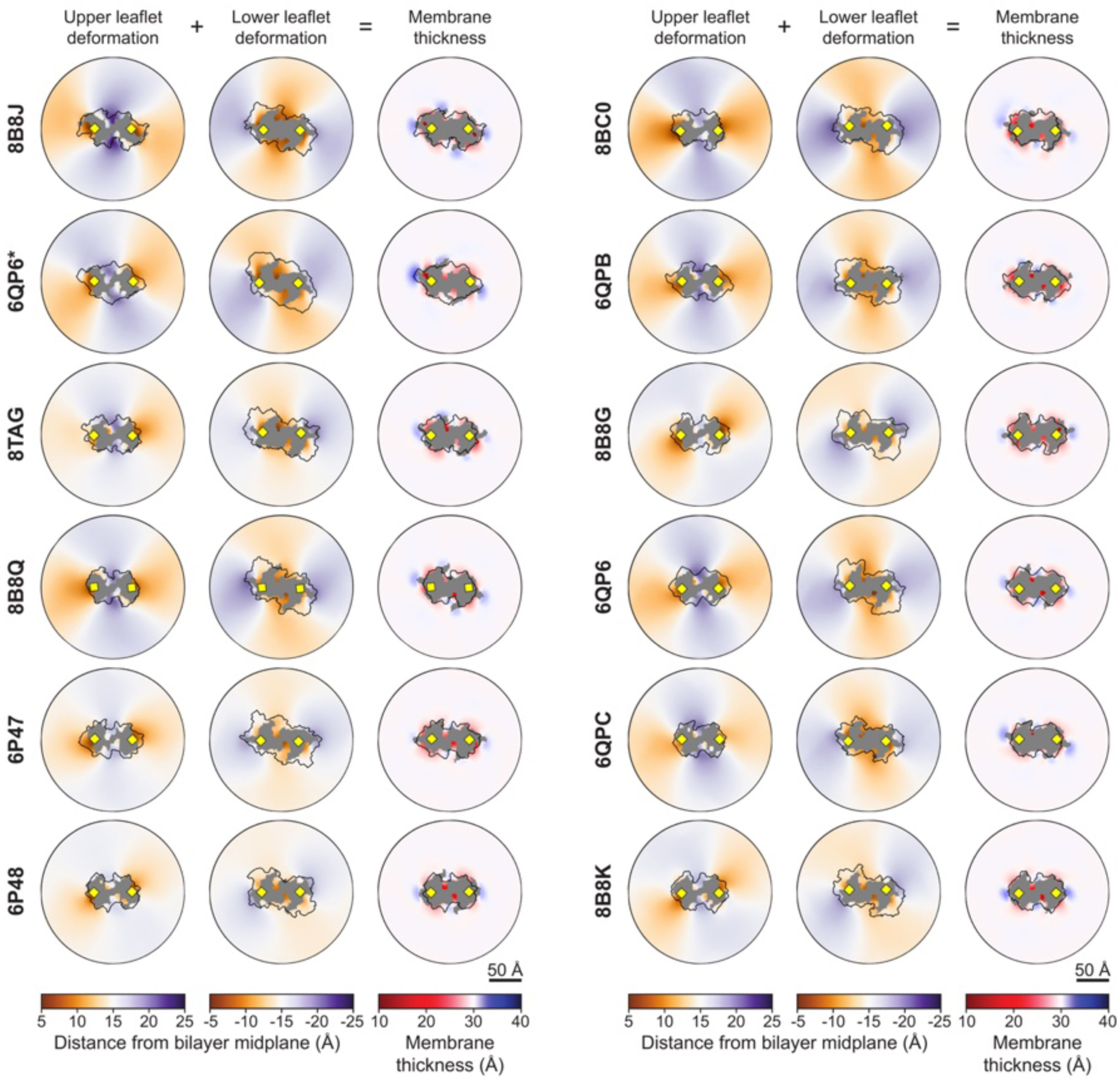
Membrane deformations for simulated TMEM16F structures. Left column: xy-map of the distance along the z-axis from the bilayer midplane to the ensemble averaged positions of the glycerol linker (GL1 and GL2 beads). Middle column: xy-map of the distance along the z-axis from the bilayer midplane to the ensemble averaged positions of the glycerol linker (GL1 and GL2 beads). Right column: the sum of the upper and lower leaflet deformations, representing the bilayer thickness along z. In all plots, grey areas indicate grid points with lipid occupancy <2%. The black outline is the projected surface of the upper (z>0) or lower (z<0) portion of the protein dimer.

**Figure 3-figure supplement 5.**
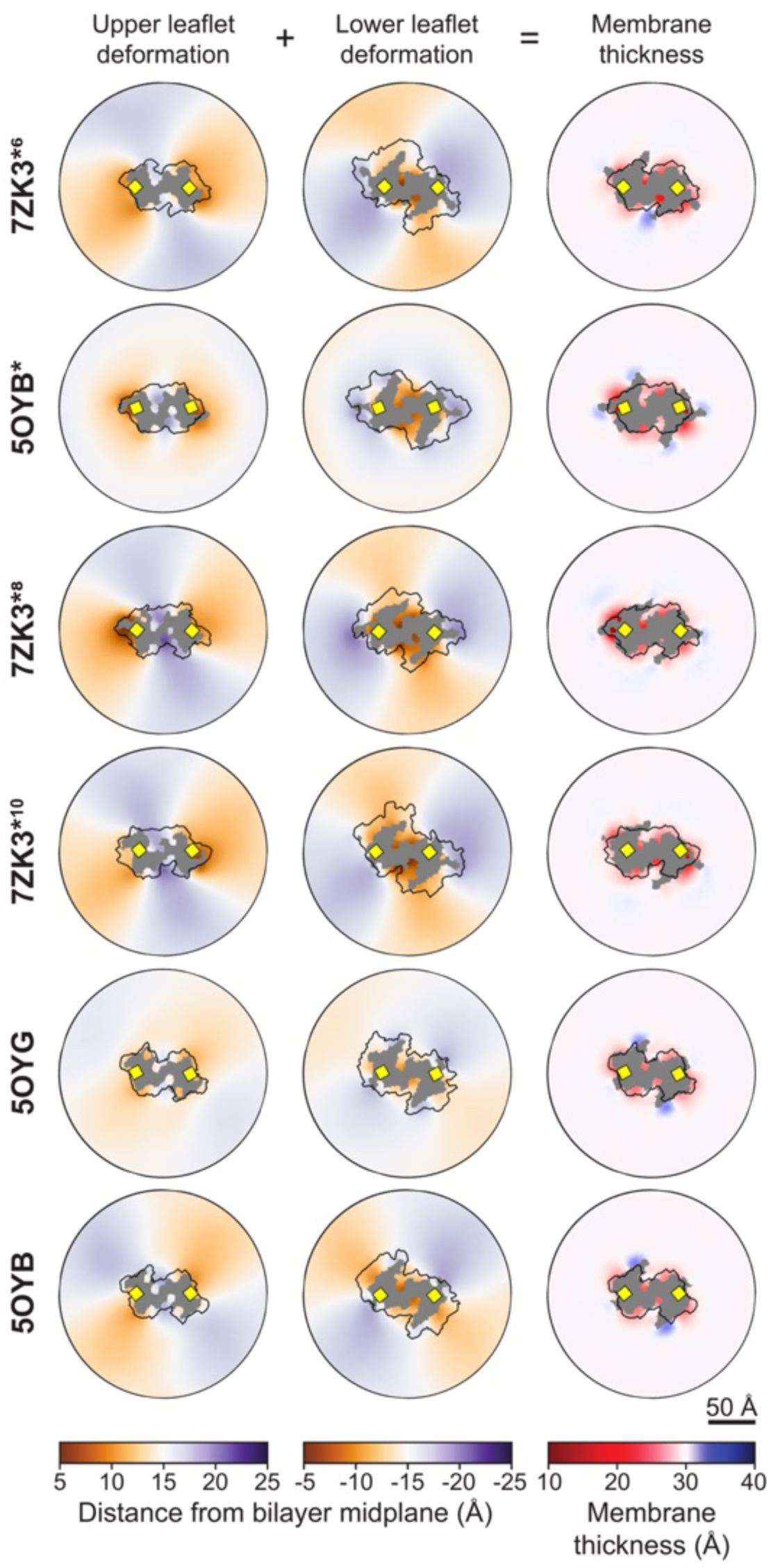
Membrane deformations for simulated TMEM16A structures. Left column: xy-map of the distance along the z-axis from the bilayer midplane to the ensemble averaged positions of the glycerol linker (GL1 and GL2 beads). Middle column: xy-map of the distance along the z-axis from the bilayer midplane to the ensemble averaged positions of the glycerol linker (GL1 and GL2 beads). Right column: the sum of the upper and lower leaflet deformations, representing the bilayer thickness along z. In all plots, grey areas indicate grid points with lipid occupancy <2%. The black outline is the projected surface of the upper (z>0) or lower (z<0) portion of the protein dimer.

**Figure 3-figure supplement 6.**
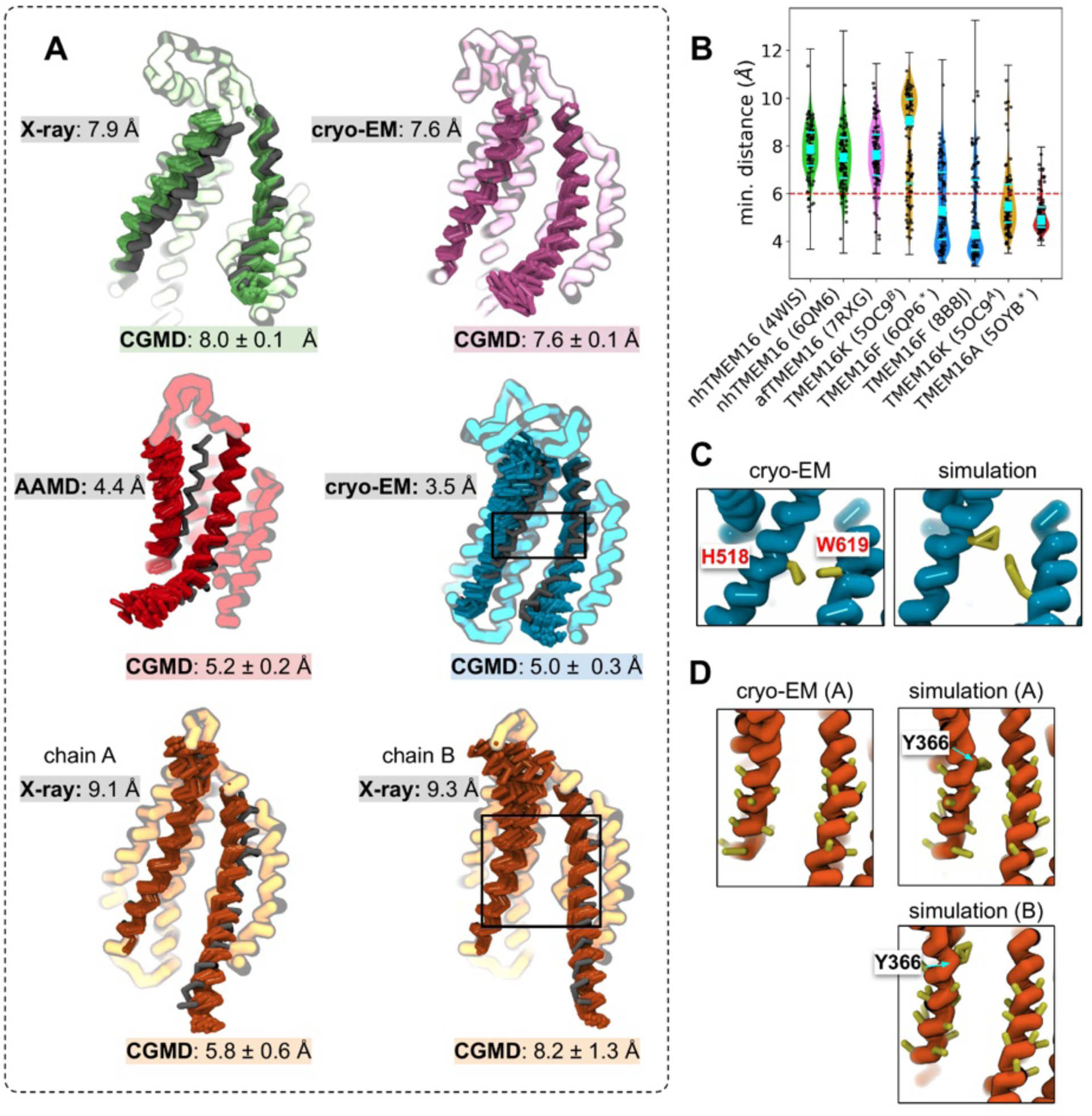
TM4 moves away from starting structure coordinates in open states. **(A)** Overlayed CG representations of experimentally determined or simulated starting structures (grey) and snapshots from CG simulations. nhTMEM16 4WIS (green), afTMEM16 7RXG (violet), TMEM16K 5OC9 (orange), TMEM16F 8B8J (blue), and TMEM16A 5OYB* (red) with minimum TM4-6 distances for the starting CG structure (grey) and mean values with standard deviation from simulations. **(B)** Violin plots of minimum distances between TM4 and TM6 in a single groove with median value (cyan square) and 25-75% quartiles (cyan bars). **(C)** TMEM16F TM4/TM6 groove constriction point at residues F518H (on TM4) and W619 (on TM6). **(D)** TMEM16K TM4/TM6 groove constriction point at Y366 (on TM4).

***Figure 3-video 1. Lipid scrambling by open Ca^2+^-bound nhTMEM16 (PDB ID 4WIS).*** *Stick representation of TM3-8 (green, rest of protein not shown) and lipid headgroups (red and blue spheres) within 7 Å of the protein. Video shows ∼600 ns of the last 1 μs of simulation with positional averaging over 3 frames. Residues T333, L336 and Y439 shown as spheres colored by residue type: hydrophobic (white) and polar (green).*

***Figure 3-video 2. Lipid scrambling by open Ca^2+^-bound TMEM16K chain B (PDB ID 5OC9).*** *Stick representation of TM3-8 (orange, rest of protein not shown) and lipid headgroups (red and blue spheres) within 7 Å of the protein. Video shows ∼600 ns of the last 1 μs of simulation with positional averaging over 3 frames. Residues Y366, I370, T435, L436 and T439 shown as spheres colored by residue type: hydrophobic (white) and polar (green).*

***Figure 3-video 3. Lipid scrambling by open Ca^2+^-bound TMEM16F F518H (PDB ID 8B8J).*** *Stick representation of TM3-8 (cyan, rest of protein not shown) and lipid headgroups (red and blue spheres) within 7 Å of the protein. Video shows ∼600 ns of the last 1 μs of simulation with positional averaging over 3 frames. Residues H518, M522 and W619 shown as spheres colored by residue type: hydrophobic (white), and polar (green).*

***Figure 3-video 4. Lipid scrambling event 1/2 by simulated ion conductive Ca^2+^-bound TMEM16A (5OYB*).*** *Stick representation of TM3-8 (pink, rest of protein not shown) and lipid headgroups (red and blue beads) within 7 Å of the protein. Scrambling lipid tail colored yellow. Residues I550, I551, K645, Q649 and E633 shown as spheres colored by residue type: acidic (red), basic (blue), polar (green) and hydrophobic (white). Video shows 3900-4780 ns of simulation with positional averaging over 3 frames.*

***Figure 3-video 5. Lipid scrambling by simulated open Ca^2+^-bound TMEM16F (PDB ID 6QP6*).*** *Stick representation of TM3-8 (cyan, rest of protein not shown) and lipid headgroups (red and blue spheres) within 7 Å of the protein. Video shows ∼600 ns of the last 1 μs of simulation with positional averaging over 3 frames. Residues F518, M522 and W619 shown as spheres colored by residue type: hydrophobic (white).*

***Figure 3-video 6. Lipid scrambling by open Ca^2+^-bound TMEM16K chain A (PDB ID 5OC9).*** *Stick representation of TM3-8 (orange, rest of protein not shown) and lipid headgroups (red and blue spheres) within 7 Å of the protein. Video shows ∼600 ns of the last 1 μs of simulation with positional averaging over 3 frames. Residues Y366, I370, T435, L436 and T439 shown as spheres colored by residue type: hydrophobic (white) and polar (green).*

**Figure 4-figure supplement 1.**
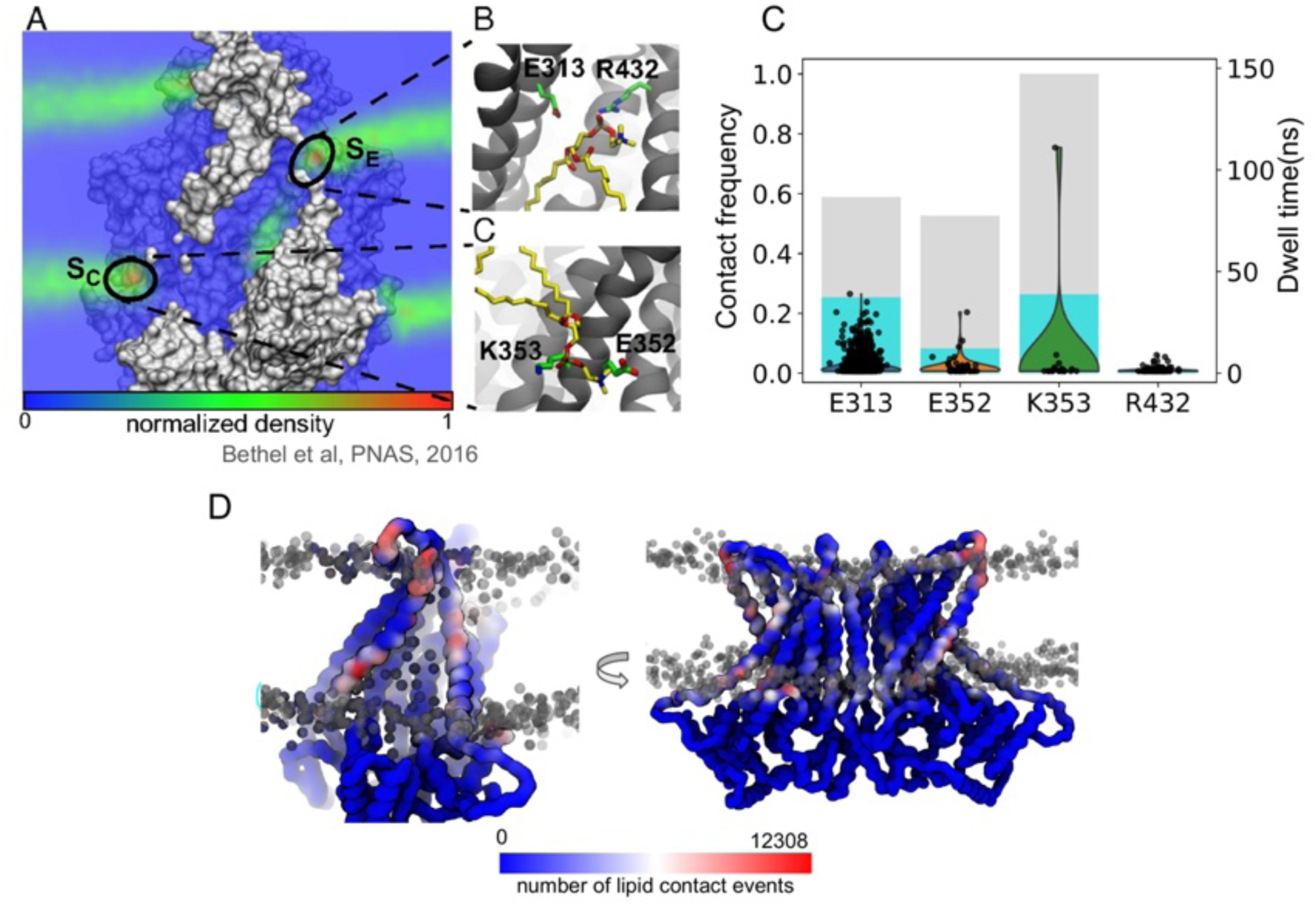
Contact analysis of lipid headgroup high density sites identified from previous AA simulation. **(A-C)** Charged residues and nearby POPC lipids near two high lipid phosphate density sites at the intracellular and extracellular entry of the open nhTMEM16 (PDB ID 4WIS) canonical groove (“S_C_”, “S_E_”) identified from previous AA simulation © 2016, Bethel & Grabe, published by PNAS (36). **(C)** Contact frequency with any lipid (grey bars) and only scrambling lipids (cyan bars). Dwell times with scrambling lipids shown as black points. **(D)** The nhTMEM16 CG backbone colored by total number of contact events with any lipid. Grey spheres indicate lipid headgroup positions in a single snapshot.

**Figure 4-figure supplement 2.**
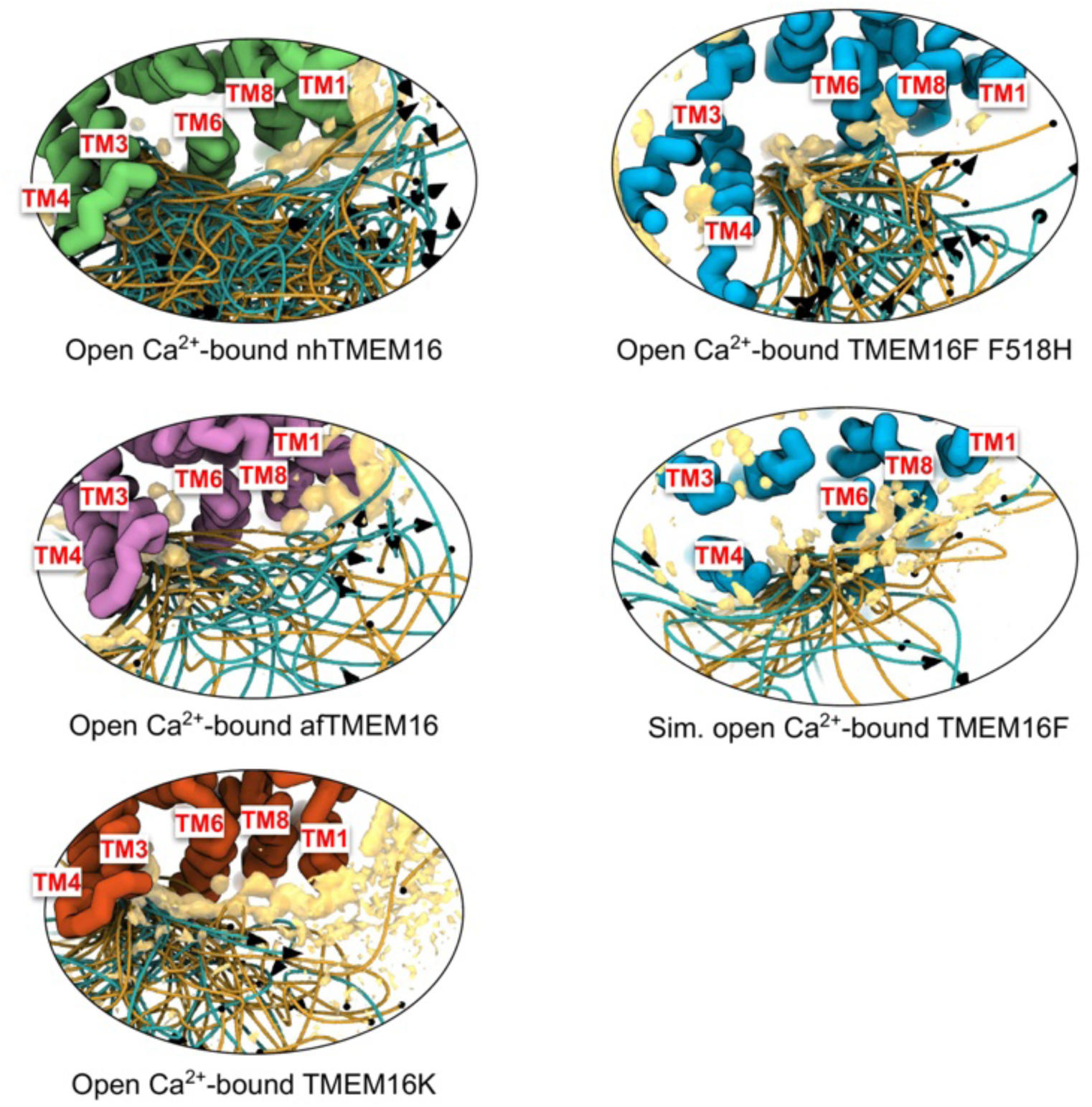
Position traces of scrambling lipids with total lipid headgroup density. Lipid traces and lipid headgroup density (yellow) for Ca^2+^ -bound scrambling competent TMEM16s: nhTMEM16 (PDB ID 4WIS, green), afTMEM16 (PDB ID 7RXG, purple), TMEM16K (PDB ID 5OC9, orange), TMEM16F F518H (PDB ID 8B8J, blue) and simulated TMEM16F (6QP6*). Each image is viewed from the extracellular or cytosolic (TMEM16K) space. Lipid traces are generated by fitting raw lipid headgroup center of mass positions to a smooth spline curve.

**Figure 4-figure supplement 3.**
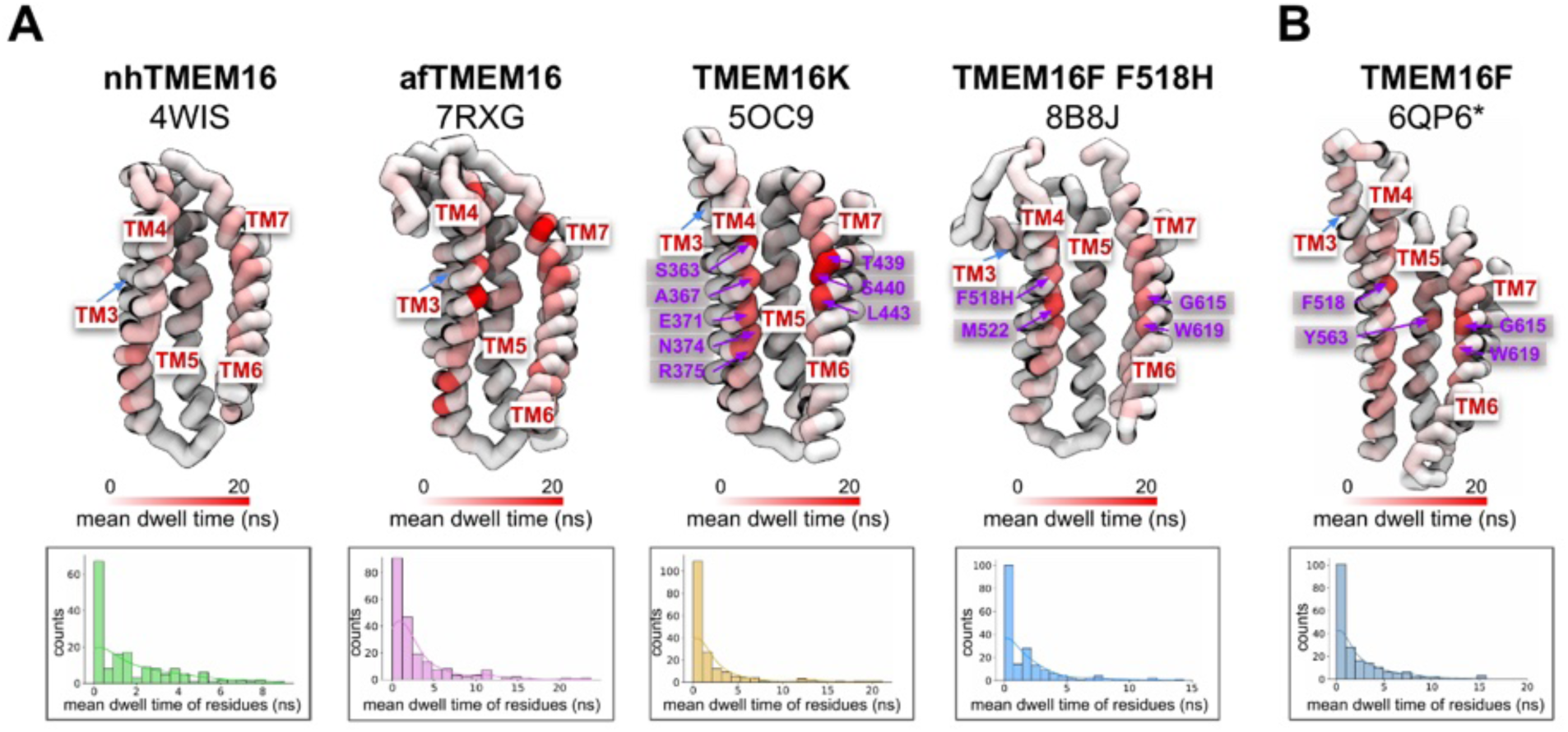
Average duration of interaction between scrambling lipids and TM4/TM6 groove lining residues. The canonical groove of each experimentally solved **(A)** and simulated **(B)** open scramblase structure is colored by the average duration of each interaction (dwell time) between scrambling lipids and groove lining residues. The distribution of average dwell times at individual residues is shown as a histogram below each structure.

**Figure 4-figure supplement 4.**
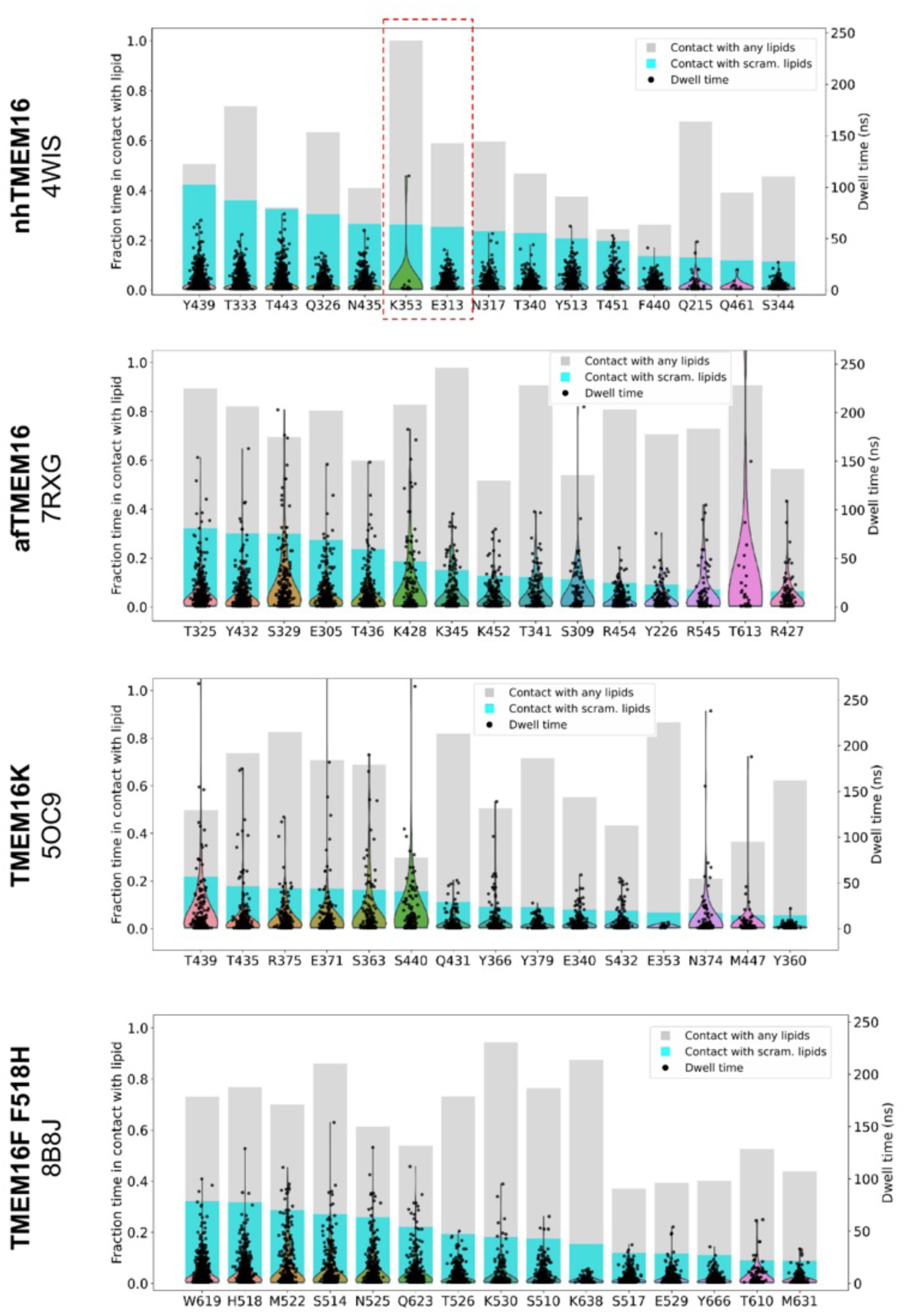
Dwell time distribution and contact frequency for TM4/TM6 groove lining residue across homologs. Contact frequency (cyan bar, left y-axis) and distribution of interaction dwell times (black scatter dots, right y-axis) between scrambling lipids and canonical groove lining residues with the 15 longest average interaction dwell times. Residues are sorted by the contact frequency. Frequency of contact with any lipid (scrambling and non-scrambling lipids taken together) is shown as grey bar. The red dashed-line rectangle indicates two previously identified residues near high lipid phosphate density in an all-atom (AA) simulation (36).

**Figure 4-figure supplement 5.**
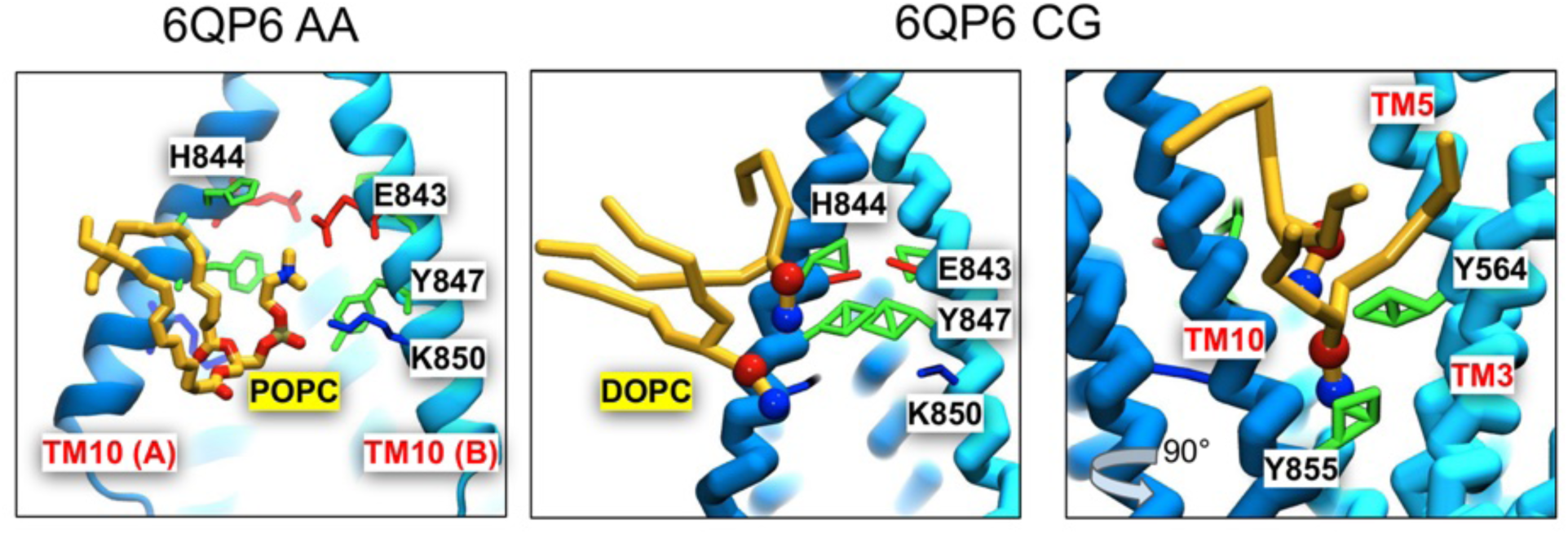
Lipids enter the dimer interface in atomistic and CG simulations of TMEM16F. Left: a snapshot of a 1-palmitoyl-2-oleoyl-glycero-3-phosphocholine (POPC) lipid that entered the TMEM16F (PDB ID 6QP6) dimer interface from the outer leaflet during an all-atom (AA) simulation. Right: snapshots at the same timepoint of 1,2-dioleoyl-sn-glycero-3-phosphocholine (DOPC) lipids that entered the TMEM16F (PDB ID 6QM6) dimer interface during a CG simulation. Nearby side chains are colored by residue type: basic (red), acidic (blue), and polar (green).

**Figure 4-figure supplement 6.**
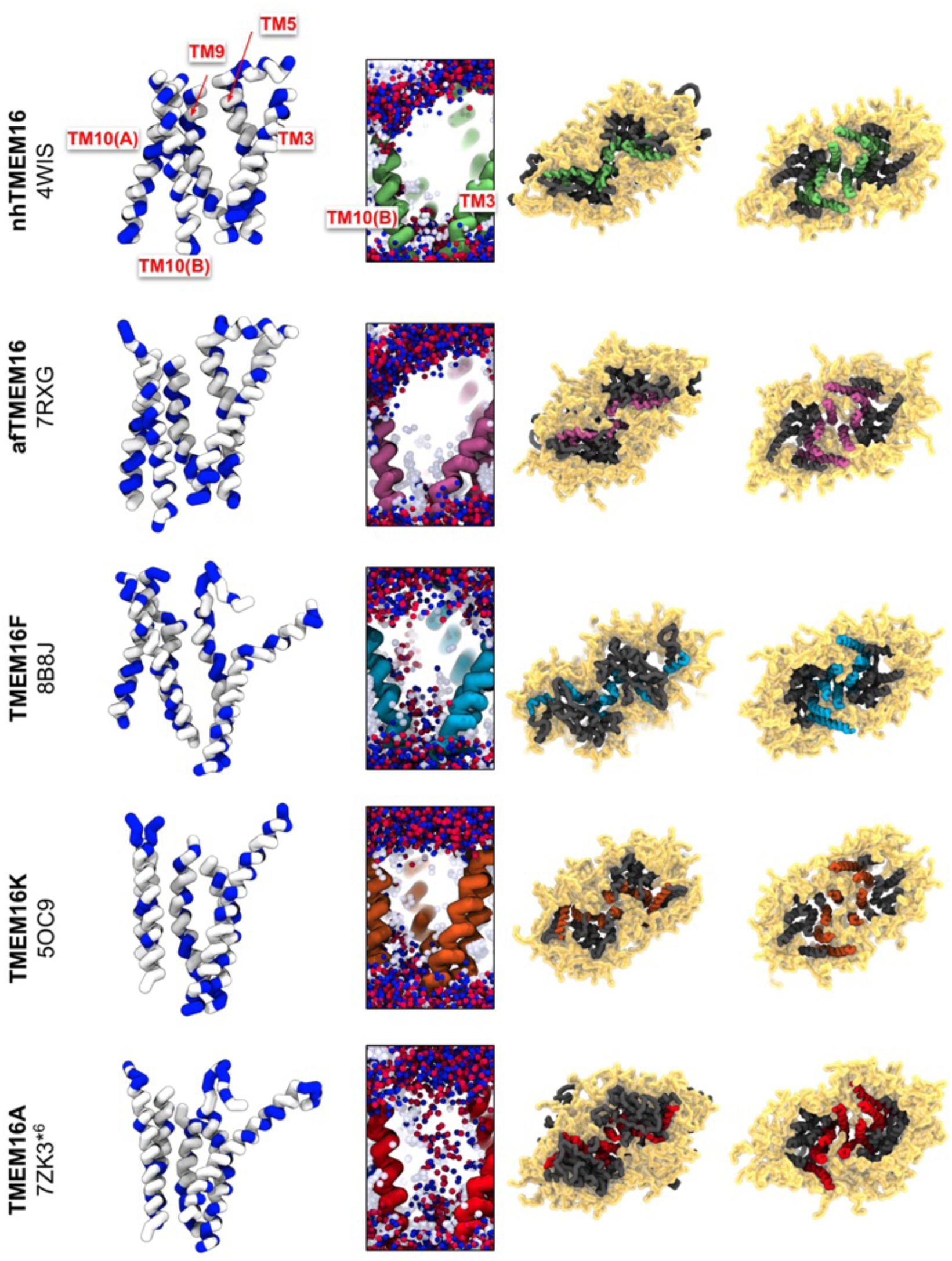
Dimer interface hydrophobicity and lipid positions. Images in each row were taken from CG MD simulations of 5 different TMEM16s. The first column depicts TM helices forming one half of the dimer interface (rest of the protein not shown) colored by residue type (small/hydrophobic: white, charge/polar: blue). The second column depicts snapshots of the same dimer interface helices with overlayed positions of water (light blue spheres) and lipid head groups (red and blue spheres) every 100 frames of the last 9 μs of each simulation. The last two columns are two views of the same snapshot showing the protein with its annulus of lipids (yellow). The dimer interface-forming helices are colored green (nhTMEM16), purple (afTMEM16), blue (TMEM16F), orange (TMEM16K), and red (TMEM16A).

**Figure 4-figure supplement 7.**
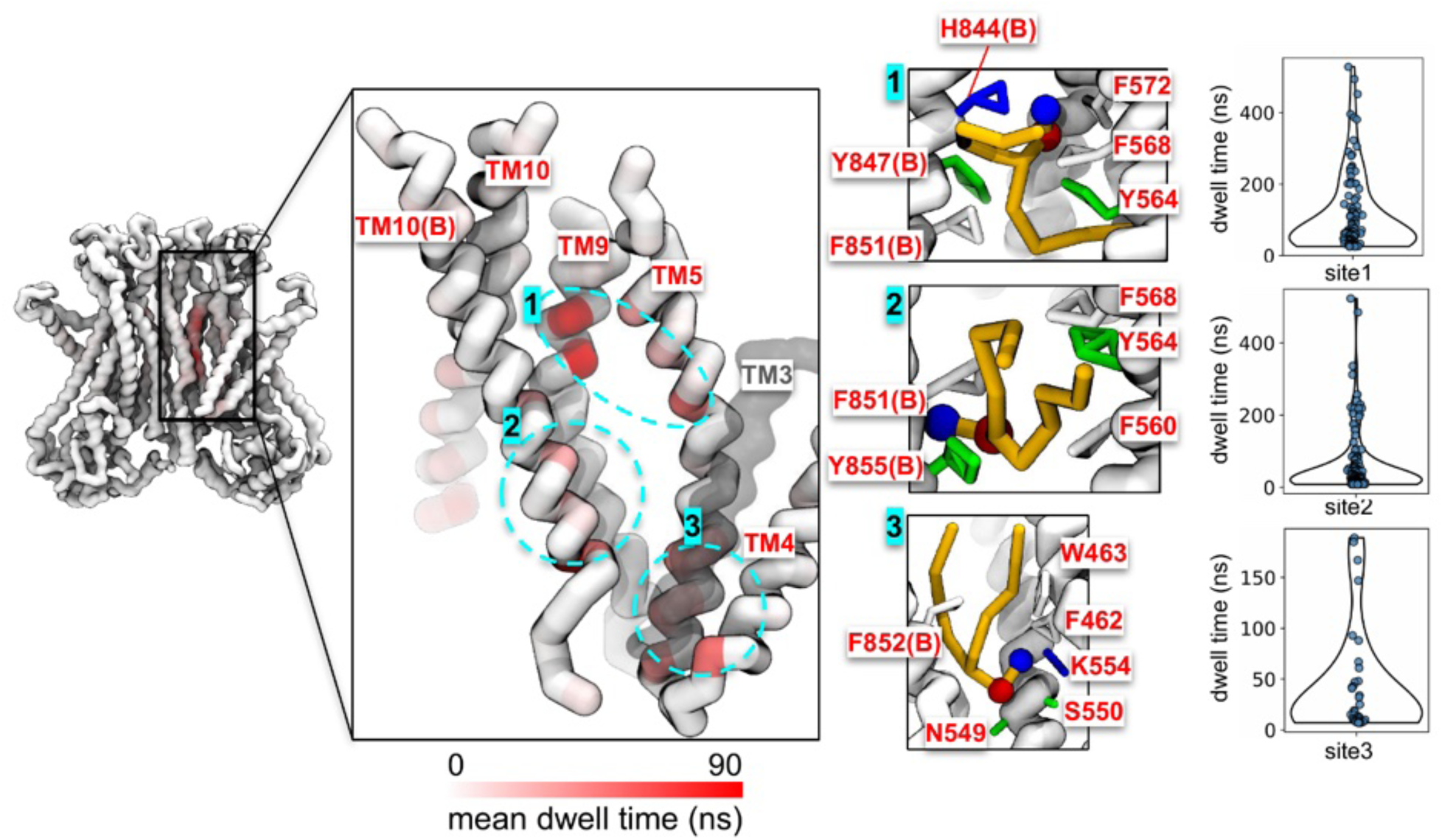
Dwell time analysis for scrambling events observed at TMEM16F F518H mutant dimer interface. The scrambling region is delineated by TM3, TM4, TM5, TM9, and TM10 both monomers. TM3 and part of TM5 are transparent for clarity. The backbone region of the dimer interface is colored by dwell time of the scrambling lipids. Sites with prolonged dwell time are circled in cyan (center left) and shown in zoomed-in images (center right). Distributions of dwell times at each site are shown as violin plots (far right).

## Notes

### Competing Interest Statement

The authors have declared no competing interest.

### Summary of Updates

In this revision of our second manuscript, we have made minor changes to main and supplemental figures and tables to either correct an error in the original plot, improve readability, or add new analysis to better represent that data. We have not significantly changed our overall interpretation of the data which remains that the groove dilation is the major determinant for scramblase activity in TMEM16s. We have added a small graphic to Figure 1A to indicate the location of the Ca2+ binding site and have added labels A through D to subfigures in Figure 4. In Figure 1, figure supplement 1 we have replaced the single snapshots from our simulations showing instantaneous membrane shapes near the protein with new images of lipid density. We think this average surface is a better comparison to the membrane deformation captured by CryoEM. In Figure 2, figure supplement 3A we corrected a label for the robustly scrambling competent subunit in asymmetric TMEM16K which was erroneously written as chain A. We have also added information about the presence of activators/modulators, Ca2+ concentration and lipid/detergent compositions uses during experimental structure determination for all structures we included in this publication (Appendix 1, Table 1). Lastly, we have added a table (Figure1, table supplement 1) to display the average number of lipid headgroups in the groove of TMEM16s with open, intermediate or closed conformations which highlights the low probability of a fully-lipid lined pore in the predicted ion conduction TMEM16A structure. In addition to these changes in figures and tables we adjusted parts of the main text to clarify our interpretation of the data. We have adjusted the language the results section Groove dilation is the main determinant for scrambling activity second paragraph to reflect the qualitative rather than quantitative relationship between membrane thinning, groove dilation and scrambling rates. In the same section we have added language to describe the dynamic behavior of the groove for different family members, as displayed in Figure 3B. Finally in the second to final paragraph of the discussion we have add one sentence to emphasize that our comparison to experimental rates should also be interpreted qualitatively as we only compare the relative rates of scrambling under activating and non-activating conditions. We think these changes remove any misleading statements that indicated a more rigorous statistical analysis of the data than was performed.

