## Appendix 1 for "Simulation-based survey of TMEM16 family reveals that robust lipid scrambling requires an open groove"

### Appendix 1 – Additional Text and Figures

#### Simulation-based survey of TMEM16 family reveals that robust lipid scrambling requires an open groove

Christina A. Stephens[†, ‡, §, 1]
Niek van Hilten[†, §, 1]
Lisa Zheng[†, ‡, §, 1]
Michael Grabe[†, #, *]

[†]Cardiovascular Research Institute
[‡]Graduate Group in Biophysics
[§]Department of Pharmaceutical Chemistry
University of California, San Francisco, CA 94158

[1] These authors contributed equally

#### Methods

##### Additional System Preparation and Simulation Details

**Starting structure selection for coarse-grained (CG) simulations**. We simulated 23 out of the 62 cryo-EM and X-ray TMEM16 homodimer structures available at the time (**Appendix 1-Table 1**). We chose not to include structures with greater than 4 Å global resolution (apart from TMEM16A PDB ID 5OYG which is the only TMEM16A apo representative) and structures with either large terminal truncations (PDB IDs 6BGI, 6BGJ, 8BC1), or largely unmodeled C-terminal domains (PDB ID 6QPI). We further narrowed our final set of structures by selecting the higher resolution of structures sharing similar backbone conformations and have the same number of $Ca^{2+}$ ions bound (**Appendix 1-Table 2**). One structure from TMEM16A (PDB ID 8QZC) and TMEM16F (PDB ID 6P46) were excluded because they share a similar conformation to other structures of the same homolog but differ in the number of bound Ca2+ ions at the orthosteric site and by a slight elevation of the TM6 C-terminus. In total, we selected 5 nhTMEM16, 2 afTMEM16, 2 TMEM16K, and 11 TMEM16F dimers. We also included 2 cryo-EM structures of TMEM16A. We also chose to simulate a computationally predicted scrambling competent TMEM16F structure based on PDB ID 6QP6 (55) which has a dilated groove similar to nhTMEM16, afTMEM16, and TMEM16K. We simulated several computationally predicted conductive states of TMEM16A: one based on $Ca^{2+}$-bound TMEM16A (PDB ID 5OYB (48)) and three based on simulations of 1PBC-bound TMEM16A (PDB ID 7ZK3) after removing 1PBC from the pore which have significant changes to TM3 and TM4 compared to their experimentally determined starting structures. Several new nhTMEM16 structures were recently published by Feng et al. (62) but not released until after completion of our simulation work and were therefore not included.

**Atomistic simulation details.** TMEM16A atomistic simulations were initiated from a $Ca^{2+}$/1PBC-bound structure (PDB ID 7ZK3) after removal of the inhibitor. The missing residues 260-266, 467-482, 526-527, 669-682 were built and refined using MODELLER (version 10.2 (87)), see more details above. Simulations were performed with Gromacs (version 2020.6 (93)) and the CHARMM36 (104) and CHARMM36m (c36m) (105) force fields for lipids and protein, respectively. PROPKA3 was used to check the protonation state of protein residues. E624 and D405 are both weakly protonatable at neutral pH, but likely well solvated and therefore left in their negatively charged states (106). The protein was embedded in a 155x155 $Å^2$ POPC bilayer and solvated in 150 mM KCl and the CHARMM TIP3P water model using CHARMM-GUI's Membrane Builder (107). System charges were neutralized using the same ions. During minimization, equilibration, and production distance restraints with 418.4 kJ mol-1 nm-2 force constants were applied between the $C\alpha$ atoms of residues 465 and 489, 454 and 566, 169 and 278, 126 and 176, 196 and 189, 123 and 282, 185 and 200 to stabilize the cytosolic domain. Simulations were run using a 2 fs time step in an NPT ensemble. Temperature was kept at 303.15 K using the Nosé-Hoover (108) thermostat (τT=1 ps). The pressure of the system was semi-isotropically coupled to a 1 bar reference pressure by the Berendsen (109) and Parrinello-Rahman (97) barostat (τP=5 ps, compressibility=4.5x10-5) for equilibration and production respectively. All bonds to H were constrained by the LINCS algorithm (110). Particle mesh Ewald (111)-calculated electrostatic and Van der Waals (VdW) interactions were cut off at 1.2 nm. A Verlet cut-off scheme was used for non-bonded interactions. VdW interactions were smoothly switched to zero between 1.0 and 1.2 nm. The protein with its bound Ca2+ ions, the membrane, and solvent bath were treated as separate groups for the thermostat coupling and center of mass removal. Harmonic restraints of the protein backbone, sidechains, lipids and dihedrals were applied and slowly reduced over 8 equilibration steps totaling ~32 ns. The equilibrated box size was ~143x143x148 $Å^3$. An identical simulation protocol was used for our atomistic simulations of TMEM16F (6QP6) but used Gromacs version 2018.7. Missing residues 84-88, 143-206, 225-228, 428-444, 490-505, 588-590, 641-644, and 792-794 were modeled using MODELLER (version 10.2 (87)). Simulation details for TMEM16A performed by the Chen group are detailed in ref. (48). Simulation details for atomistic simulations of TMEM16F (6QP6) performed by the Weinstein group are detailed in ref. (55). Simulation details for atomistic simulations for nhTMEM16 are detailed in ref. (36).

**Simulated TMEM16A structure selection.** TM4/TM5/TM6 residue pair distances from aggregate atomistic trajectories of TMEM16A were submitted to time-independent component analysis (tICA) (112) and subsequent K-medoids clustering. 100 clusters were then used to construct a Markov-state model and microstates were grouped into macrostates using the improved Perron-cluster cluster analysis (PCCA+) method (113). The above analysis was performed using MSMBuilder (114). We selected the medoids from three of these macrostates (cluster 6, 8, and 10) which are more dilated than the starting structure and predicted to represent different conductive states (**Appendix 1-Figure 1**).

**Additional Simulation Analysis Methods**

**Maximum density path calculations.** One-dimensional paths through the lipid densities were calculated by first selecting a single grid cell near the center of the box, totaling the density of all cells within a 4.7 Å cutoff and then repeating this step for each cell within a 9.4 Å of the first, until a maximum density total was identified. We then saved the centroid of this final set of grid cells and repeated the search using this centroid, or node, as the starting position. This process was continued until either the path length (sum of distances between subsequent nodes) reached 80 Å for lipids (30 Å for water), no new nodes were found, or the next selected node caused the path to deviate sharply (<90°). We only included cells with densities >= 0.0005 for lipids or >= 0.002 for water for this calculation. Path nodes were then interpolated using a B-spline representation in Scipy methods (100) and final nodes were selected from this path to given 0.5 Å gaps between nodes for the lipid paths and 4.7 Å for water.

**Water and ion permeation analysis.** For each simulation the positions of water and ion CG beads within a 9.4 Å search radius of the water maximum density pathway were tracked overtime using a custom script that includes MDAnalysis methods (98, 99). Water beads were assigned to path nodes if they fell within the bounds of a cylindrical disc (4.7 Å height, 9.4 Å diameter) centered on the bead with its face normal defined by the vector between the current and subsequent node. Permeation events were counted if a water or ion left the search radius into the bath opposite to the one from which it entered the path. The maximum density path was mirrored symmetrically on both subunits and each pathway was tracked independently. We also calculated the flux of water between nodes by counting the net number of waters entering and leaving a cylinder (of the same proportions above) centered on each path node at each 1 ns timestep.

**Diffusion coefficient calculations.** Diffusion coefficients for lipids were calculated by first making a 3D interpolated path for each scrambling event and then measuring the minimum path distance to the starting position at each time point. We used a linear least squared regression to fit the slope of each squared displacement curve (squared minimum path distance) and divided the slope by 2 (for 1D diffusion) to obtain the final diffusion coefficient for each lipid. We wrote custom scripts for this analysis using MDAnalysis (98, 99) and Scipy methods (100).

**PMF calculations.** To calculate the PMFs for each pathway, we assigned each cell from the average PC density to its closest respective mean-lipid pathway node. We then summed the density of all cells assigned to a given node and divided by the total volume of those cells. We used the following equation to convert the density values to energies or PMF.

$$E = \frac{-R*T*\ln(\rho/\rho_0)}{1000} \qquad (1)$$

$\rho_0$ is the density of PC in a 10 $\mu$s protein-free membrane simulation, *R* is the gas constant.

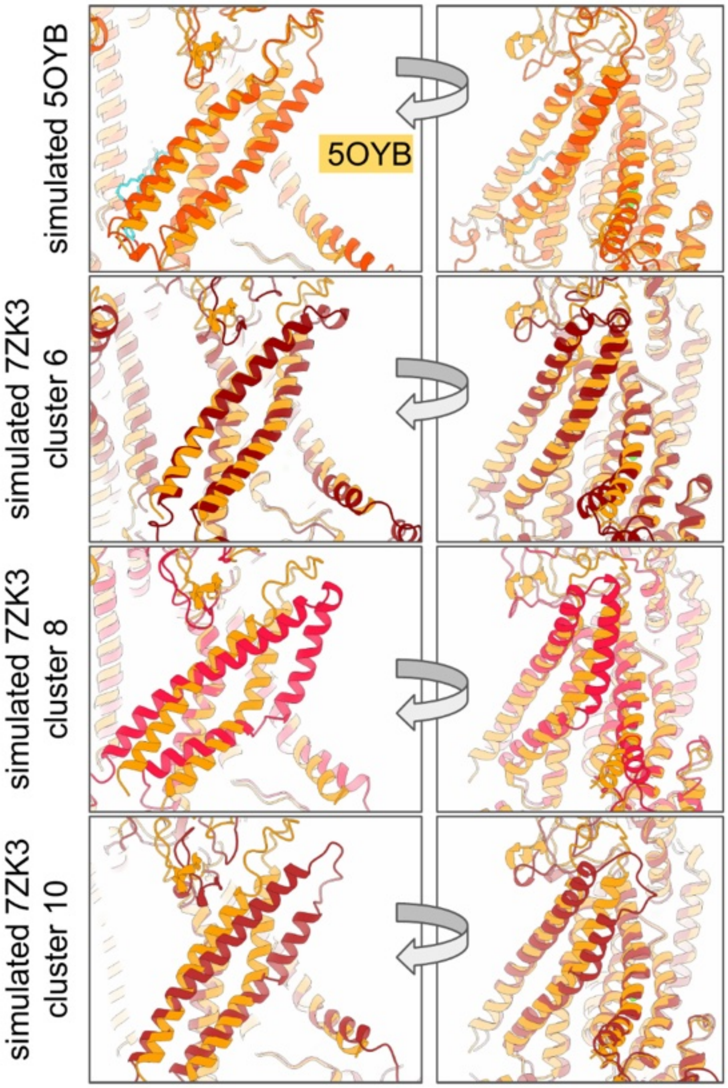

***Appendix 1-Figure 1. Predicted alternative and ion conductive states of TMEM16A.*** *Starting coordinates for simulated conductive states of TMEM16A generated from atomistic MD simulations (darker colors). Each predicted open subunit is aligned to a $Ca^{2+}$-bound TMEM16A cryo-EM structure (PDB ID 5OYB, light orange).*

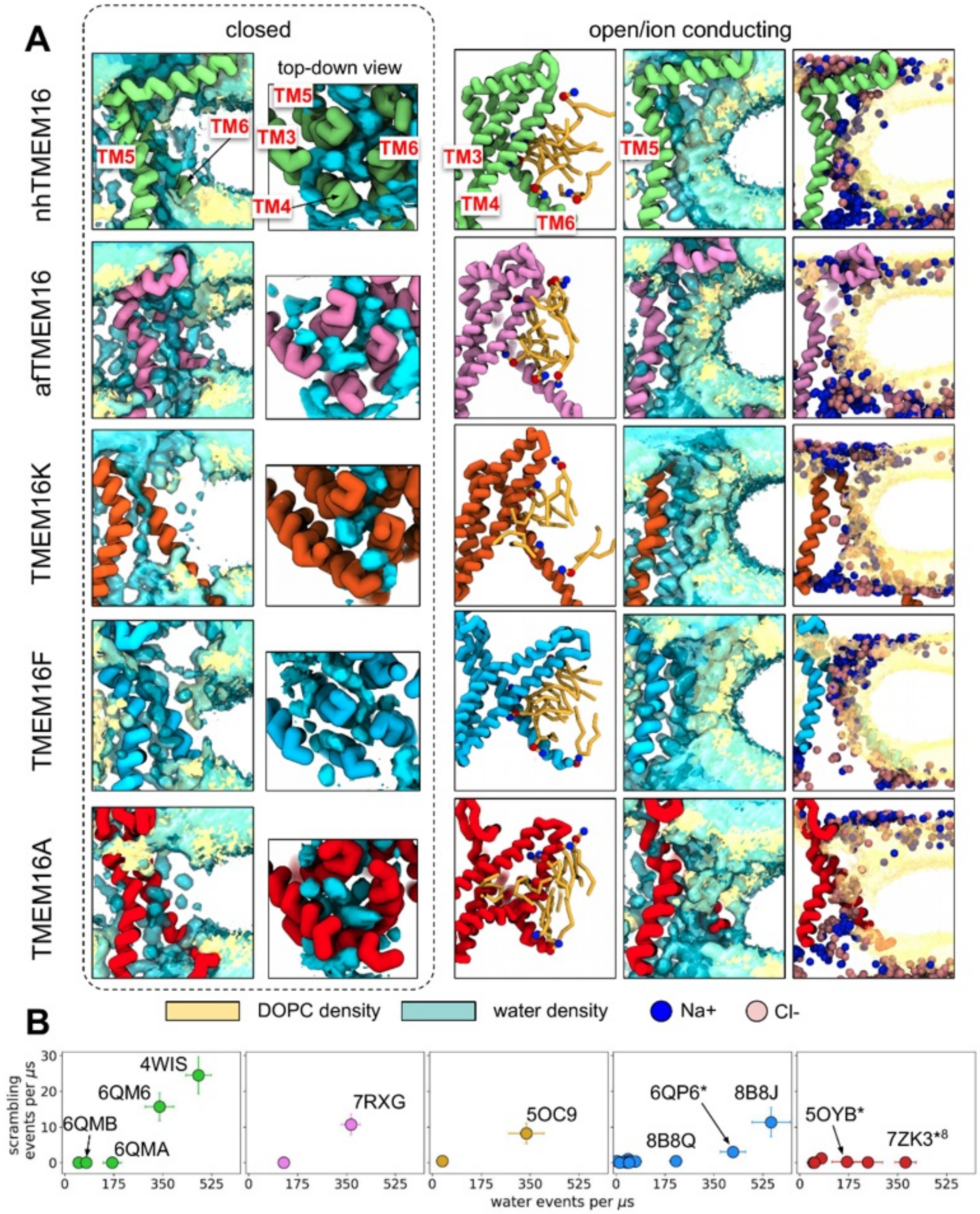

***Appendix 1-Figure 2. Simulated lipid scrambling correlates with water permeation through the TM4/TM6 groove. (A)*** *Water and lipid headgroup density in CG simulations of closed (**left**) and open/ion conducting (**right**) structures: nhTMEM16 (PDB IDs 6QM4 (closed) and 4WIS (open), green), afTMEM16 (PDB IDs 7RXB (closed) and 7RXG (open), violet), TMEM16K (PDB IDs 6R7X (closed) and 5OC9 (open), gold), TMEM16F (PDB IDs 6QPB (closed) and 8B8J (open), blue) and TMEM16A (PDB ID 5OYG and simulated 7ZK3*[8]). Ions positions (cyan and green beads) shown every 100 frames. TM4 not shown in the first and two rightmost columns. Density is shown at the same isovalue contour for all images. **(B)** The number of scrambling events plotted against the average number of water permeation events per μs. Due to the presence of asymmetric structures the maximum water permeation rate of the two subunits is shown. Water beads were tracked within a 9.4 Å radius of the maximum density pathway (see SI Methods). Density is shown at the same isovalue contour for all images.*

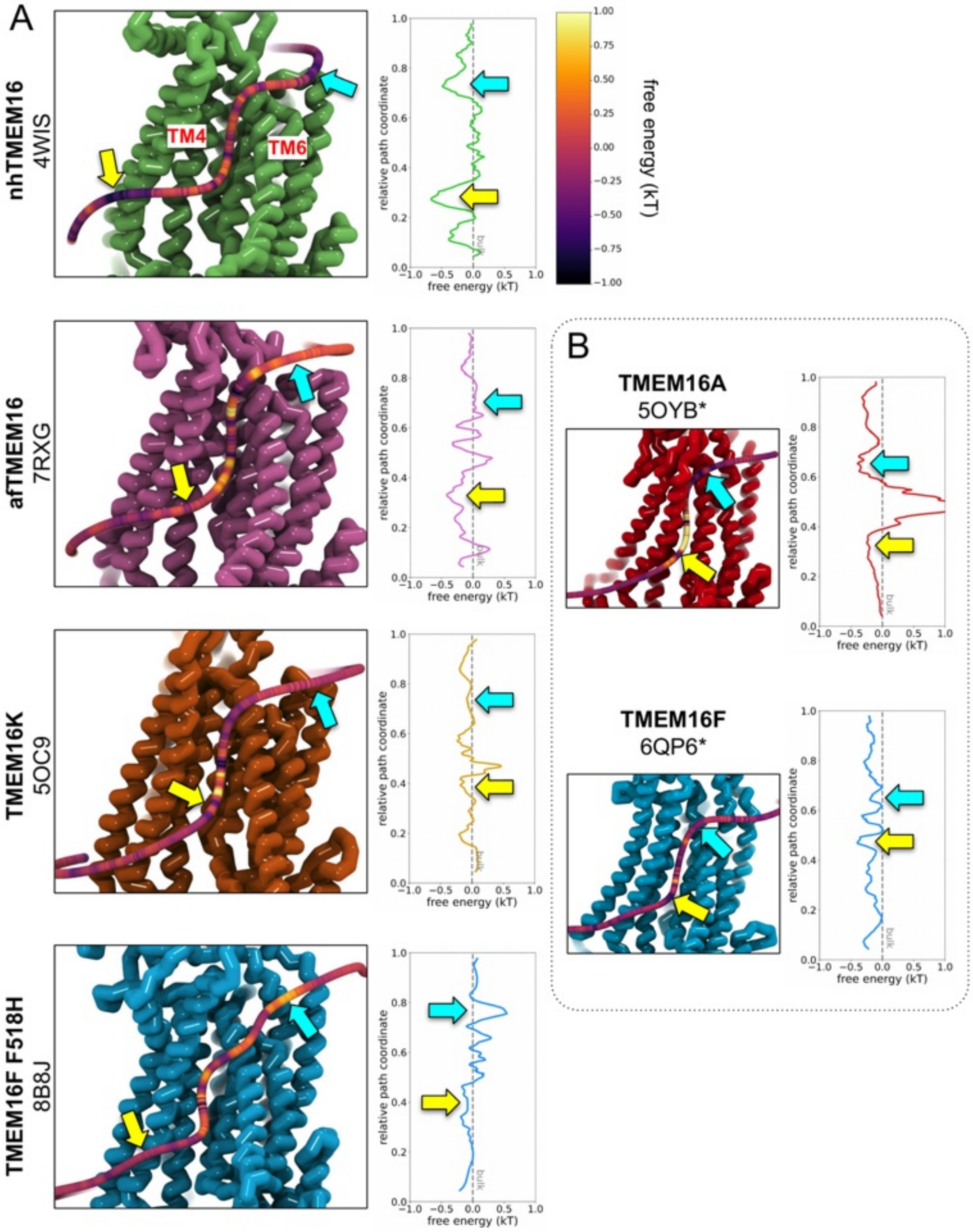

***Appendix 1-Figure 3. PMF estimates from DOPC headgroup density. (A)*** *Left: Snapshots of nhTMEM16 (PDB ID 4WIS, green), afTMEM16 (PDB ID 7RXG, violet), TMEM16K (PDB ID 5OC9, orange), and TMEM16F F518H (PDB ID 8B8J, blue), overlaid with a 3D maximum density path (see methods) calculated from DOPC headgroup positions averaged from both subunits, except for asymmetric TMEM16K and WT TMEM16F paths on subunit with higher scrambling events. Each path is colored by the free energy (kT) calculated from the total density within 4.7 Å of a given node on the path normalized by the lipid headgroup density from simulation of a protein-free DOPC bilayer. Right: the same PMF plotted against its relative position along the maximum density pathway. Colored arrows indicate the same locations along the path.* ***(B)*** *PMF-colored 3D maximum lipid density paths for simulated ion conducting TMEM16A (5OYB*, red) and a simulation open WT TMEM16F (6QP6*, blue) simulation with corresponding PMF curve (right).*

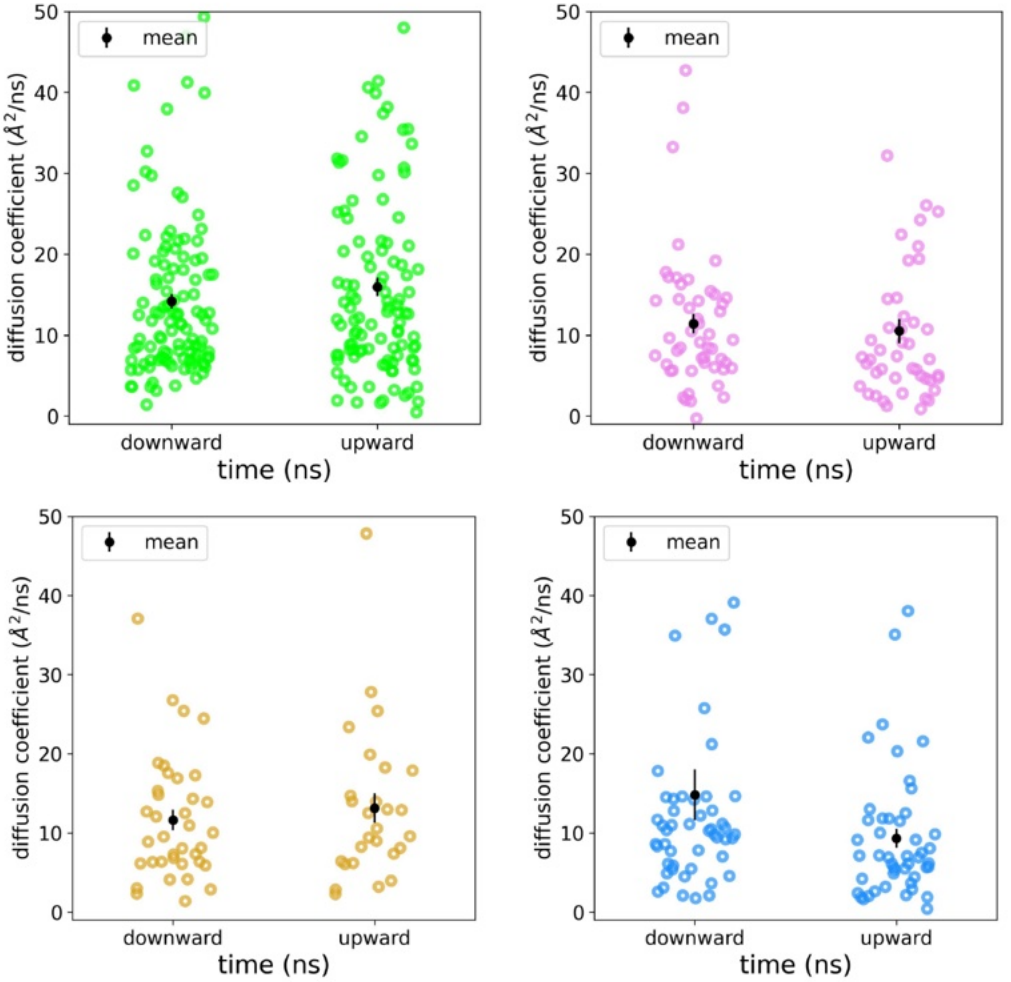

***Appendix 1-Figure 4. Diffusion coefficients for scrambling lipids.*** *Diffusion coefficients were fit to individual mean squared displacement curves for individual DOPC lipids during their scrambling event in simulations of nhTMEM16 (PDB ID 4WIS, green), afTMEM16 (PDB ID 7RXG, violet), TMEM16K (PDB ID 5OC9, gold), and TMEM16F (8B8J, blue). The mean with standard deviation shown in black.*

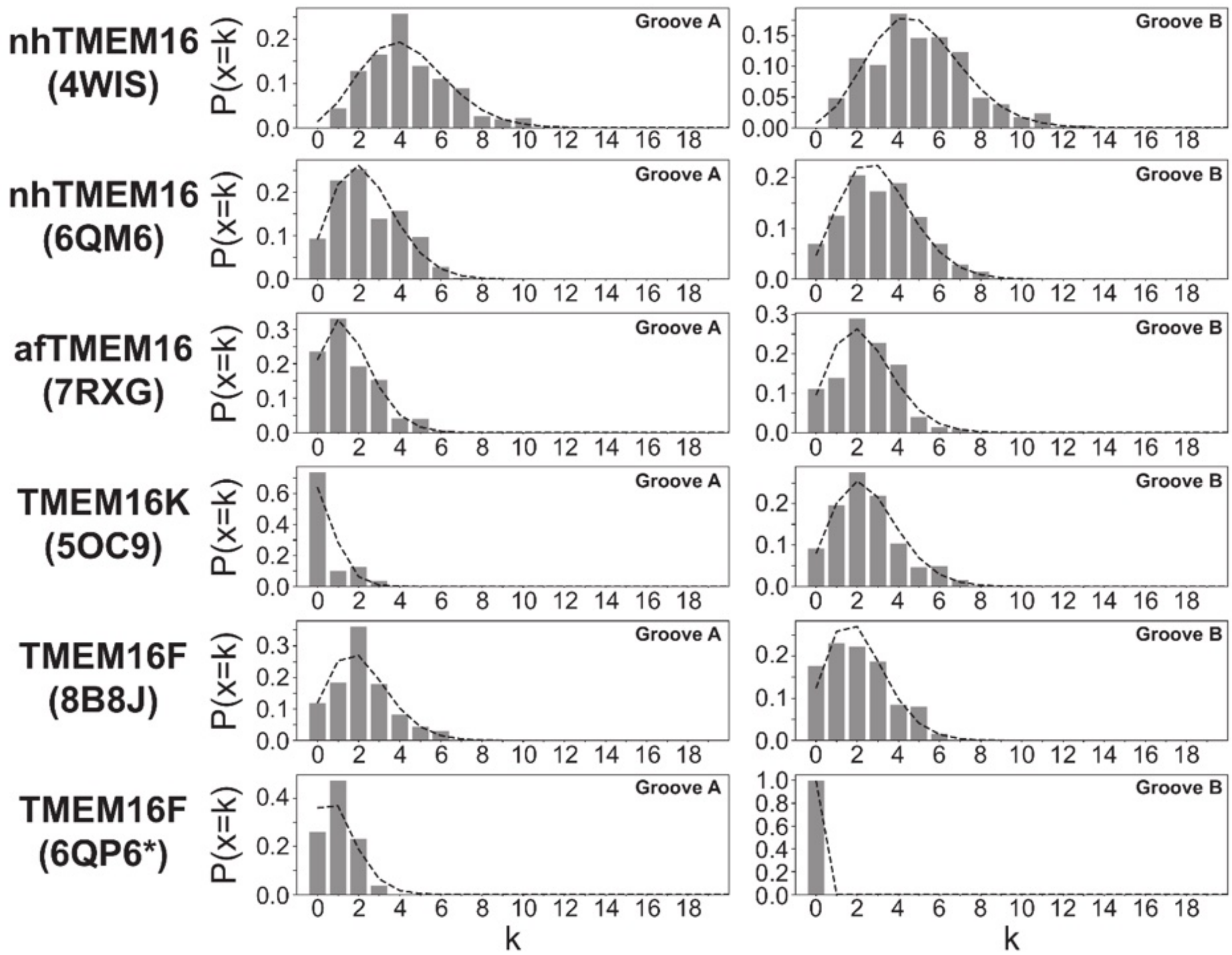

***Appendix 1-Figure 5. Scrambling events are Poisson distributed.*** *Poisson distributions calculated on 400 ns intervals for all scrambling events in groove A (left) and groove B (right) of scrambling-competent structures. Bars represent data, dashed line is the theoretical distribution* $P(x=k) = \lambda^k e^{-\lambda}/k!$*, with* $\lambda$ *being the average number of events in the 400 ns interval.*

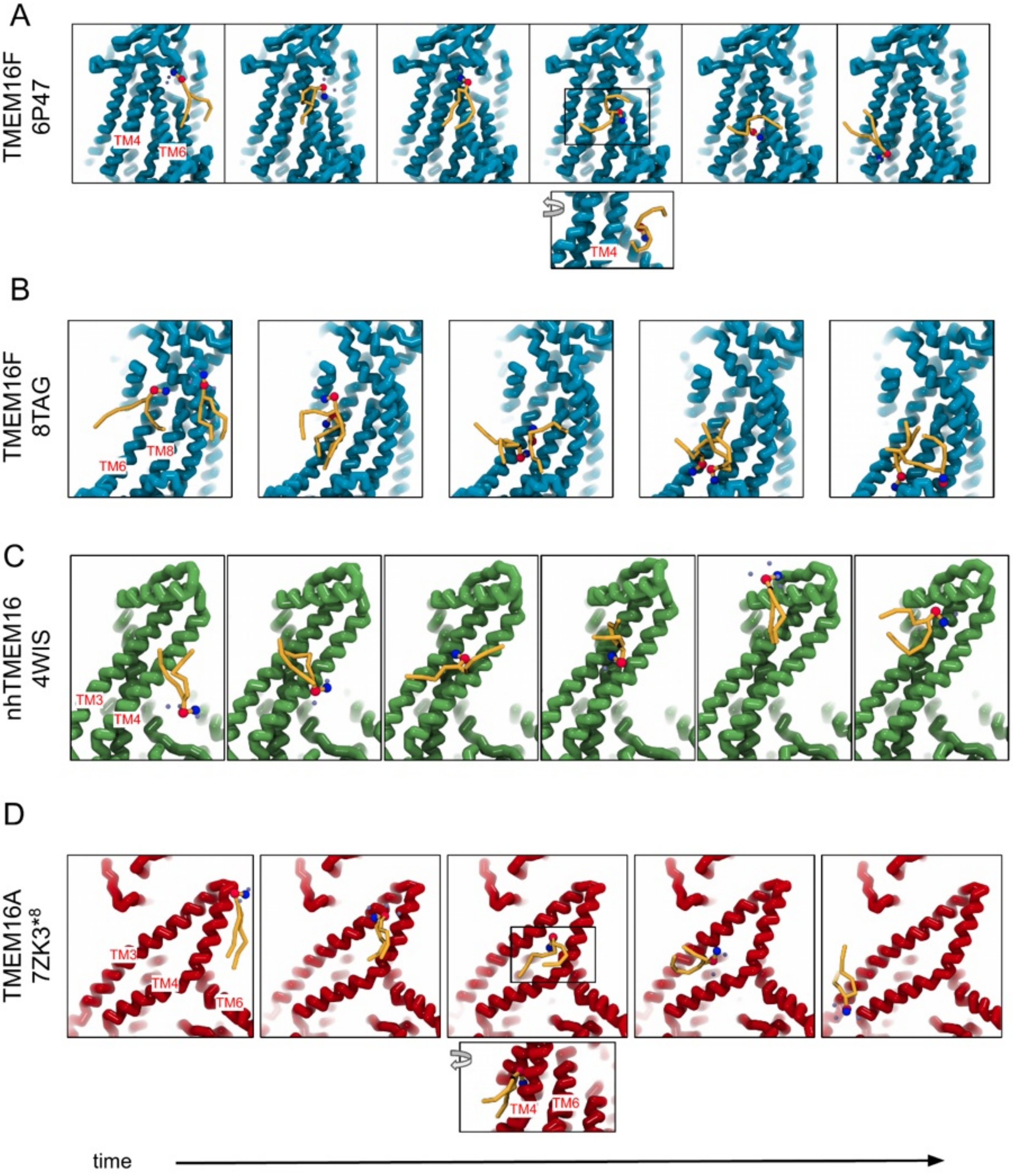

***Appendix 1-Figure 6. "Out-of-the-groove" scrambling events outside of the dimer interface.*** ***(A)*** *Snapshots over time of a single lipid scrambling event by TMEM6F over the closed TM4/TM6 groove.* ***(B)*** *Snapshots over time of two concurrent scrambling events by TMEM16F over TM6 and TM8.* ***(C)*** *Snapshots over time of a single lipid scrambling event by nhTMEM6 over TM3 and TM4.* ***(D)*** *Snapshots over time of a single lipid scrambling event by TMEM6A over TM3 and TM4.*

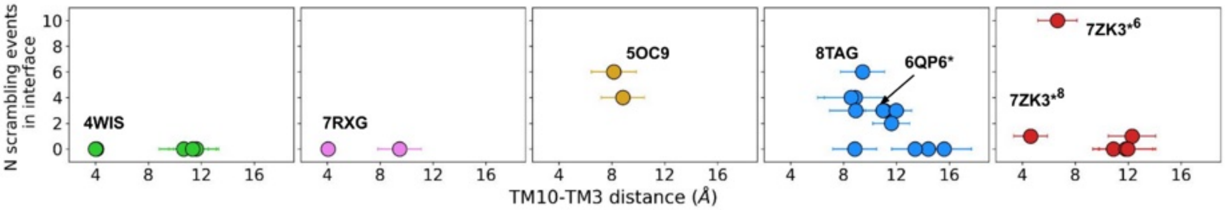

***Appendix 1-Figure 7. Number of scrambling events and the width of the dimer interface inner leaflet entrance.*** *The number of scrambling events at the dimer interface plotted against the average minimum distance between TM10 and TM3 on the opposite helix near the inner leaflet-solvent boundary.*

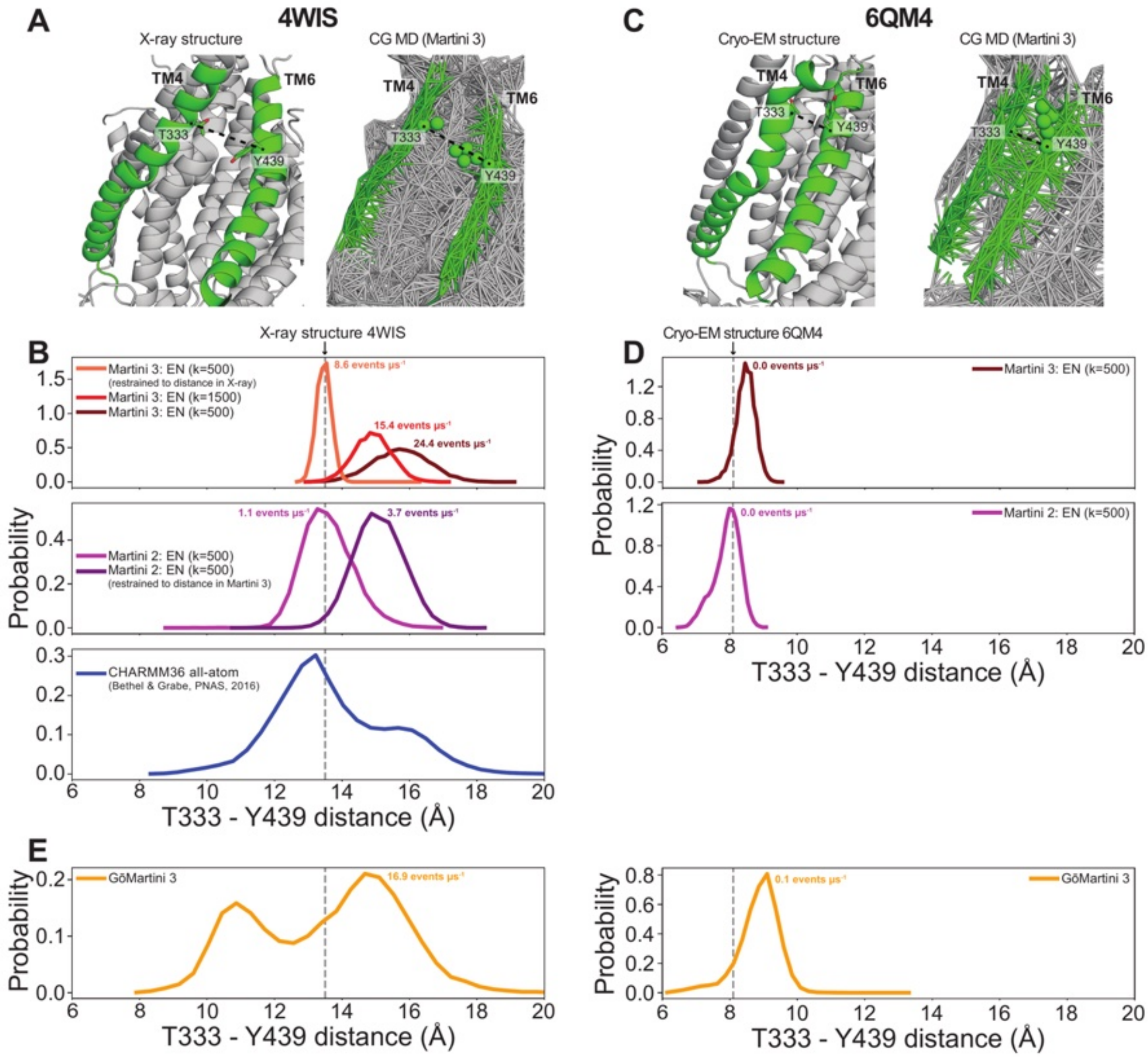

***Appendix 1-Figure 8. TM4/TM6 groove distances vary between different force fields. (A)*** *The X-ray structure (left) and the full elastic network for the last frame of our CG MD Martini 3 simulation (right) of open-groove nhTMEM16 (4WIS). TM4 and TM6 are colored green. The dashed line indicates the distance between the Cα-atoms/backbone beads of T333 and Y439, which is used to measure TM4/TM6 distance in B and E.* ***(B)*** *Probability distributions of the T333-Y439 distance in 4WIS for different settings of the Martini 3, Martini 2, and CHARMM36 force fields. EN = elastic network. k indicates the force constant used in the EN. Scrambling rates are calculated as described throughout the current work. The CHARMM36 data was taken from ref. (36). Dashed gray line indicates the distance in the X-ray structure.* ***(C)*** *The cryo-EM structure (left) and the full elastic network for the last frame of our CG MD Martini 3 simulation (right) of closed-groove nhTMEM16 (6QM4). TM4 and TM6 are colored green. The dashed line indicates the distance between the Cα-atoms/backbone beads of T333 and Y439, which is used to measure TM4/TM6 distance in D-E. Note the elastic network connections between TM4 and TM6 in green, that are absent in A.* ***(D)*** *Probability distributions of the T333-Y439 distance in 6QM4 for different settings of the Martini 3 and Martini 2 force fields. Dashed gray line indicates the distance in the X-ray structure.* ***(E)*** *Probability distributions of the T333-Y439 distance in 4WIS (left) and 6QM4 (right) for simulations with the GōMartini 3 force field (84). Dashed gray line indicates the distance in the respective experimental structures*

***Appendix 1 - Table 1. TMEM16 structures selected for molecular dynamics (MD) simulations, the concentrations they were solved/constructed in, and modeled missing residues.***

| homolog | PDB ID | structure method | detergent/lipid environment | activator/ modulator | organism | modeled loops |
|---|---|---|---|---|---|---|
| **nhTMEM16** | 4WIS | X-ray | n-dodecyl-β-D-maltopyranoside (DDM, Anatrace) | 3mM $CaCl_2$ | fusarium vanettenii 77-13-4 (fungus) | 128-140,465-482,586-593,657-659,685-691 |
| nhTMEM16 | 4WIS | X-ray + Amber/c36m | POPC | 2 $Ca^{2+}$ per chain | fusarium vanettenii 77-13-4 (fungus) | 128-140,465-482,586-593,657-659,685-691, |
| **nhTMEM16** | 6QM4 | Cryo-EM | MSP2N2 nanodisc with POPC:POPG=7:3 | No | fusarium vanettenii 77-13-4 (fungus) | 1-14,417-424,586-594,651-664,685-687 |
| **nhTMEM16** | 6QM6 | Cryo-EM | DDM (Anatrace) | No | fusarium vanettenii 77-13-4 (fungus) | 1-14,416-418,468-476,588-594,653-663,685-687 |
| **nhTMEM16** | 6QMA | Cryo-EM | MSP2N2 nanodisc with POPC:POPG=7:3 | 0.3mM $CaCl_2$ | fusarium vanettenii 77-13-4 (fungus) | 1-14,417-424,469-476,586-594,651-664,685-687 |
| **nhTMEM16** | 6QMB | Cryo-EM | MSP2N2 nanodisc with POPC:POPG=7:3 | 0.3mM $CaCl_2$ | fusarium vanettenii 77-13-4 (fungus) | 1-14,417-424,468-476,586-594,651-664,685-687 |
| **afTMEM16** | 7RXB | Cryo-EM | nanodisc with DOPC:DOPG=7:3 | No | Aspergillus fumigatus (fungus) | 1-10,25-31,90-102,256-268,312-317,443-463, 594-603 |
| **afTMEM16** | 7RXG | Cryo-EM | MSP1E3 nanodisc with DOPC:DOPG=7:3 | 0.5mM $CaCl_2$ | Aspergillus fumigatus (fungus) | 1-11,725-735 |
| **TMEM16K** | 5OC9 | X-ray, LCP | 1-(7Z-hexadecenoyl)-rac-glycerol | 0.1M $CaCl_2$ | Homo sapiens (mammal) | ASYMMETRIC 1-13A,57-66A,188-191A,350-351A,472-474A,1-13B,188-191B,348-353B |
| **TMEM16K** | 6R7X | Cryo-EM | undecyl maltoside and chloestryl hemisuccinate | 2mM $CaCl_2$ | Homo sapiens (mammal) | 1-12 |
| **TMEM16F** | 6P47 | Cryo-EM | digitonin | 0.5mM $CaCl_2$ | Mus musculus (mammal) | 82-88,197-203,257-258,431-444,490-498,639-649 |
| **TMEM16F** | 6P48 | Cryo-EM | MSP2N2 nanodisc with POPC:POPE:POPS=3:1:1 | 2mM $Ca^{2+}$, $PIP_2$ (1 $PIP_2$ per 50 lipid molecules) | Mus musculus (mammal) | 77-89,131-133,225-231 |
| **TMEM16F** | 6QPB | Cryo-EM | digitonin | No | Mus musculus (mammal) | 80-89,491-501,633-645,791-792 |
| **TMEM16F** | 6QP6 | Cryo-EM | digitonin | 1mM $CaCl_2$ | Mus musculus (mammal) | 82-86,197-201,224-227,489-502,588-590,641-644,792-794 |
| **TMEM16F** | 6QPC | Cryo-EM | MSP2N2 with POPC:POPG=3:1 | 1mM $CaCl_2$ | Mus musculus (mammal) | 108-114,258-263,639-646,765-771 |
| **TMEM16F** | 8TAG | Cryo-EM | MSP2N2-SoyPC nanodisc | 4mM $CaCl_2$, $PIP_2$ ($PIP_2$:TMEM16 monomer=4:1) | Mus musculus (mammal) | ASYMMETRIC 52-58A, 83-90A, 223-231A, 639-644A, 490-502B, 869-871B |
| **TMEM16F** | 8B8G | Cryo-EM | digitonin | No | Mus musculus (mammal) | 80-89,491-501,637-645,790-794 |
| **TMEM16F** | 8B8J | Cryo-EM | digitonin | 2mM $CaCl_2$, 0.01mM diC8-$PI_{(4,5)}P_2$ | Mus musculus (mammal) | 82-88,120,221-232,489-502,641-644,792-794 |
| **TMEM16F** | 8B8Q | Cryo-EM | MSP2N2 nanodisc with POPC:POPG=3:1 | 2mM $CaCl_2$ | Mus musculus (mammal) | 82-88,120,221-232,639-644,789-794 |
| **TMEM16F** | 8B8K | Cryo-EM | digitonin | 2mM $CaCl_2$, 0.01mM diC8-$PI_{(4,5)}P_2$ | Mus musculus (mammal) | 82-86,197-201,224-227,489-502,588-590,641-644,792-794 |
| **TMEM16F** | 8BC0 | Cryo-EM | glyo-dendrimer (GDN, Anatrace) | 2mM $CaCl_2$ | Mus musculus (mammal) | ASYMMETRIC 79-90A, 220-232A,254-263A,588-590A,640-644A,792-794A, 79-90B,120B,220-232B,254-263B,639-644B |
| TMEM16F | 6QP6 | Cryo-EM + OpenMM+c36m | POPE:POPG=7:3 | 1 Ca2+ per chain, plus 1 Ca2+ at dimer interface | Mus musculus (mammal) | 1–42, 150–186, 428–444, 489–502, 876–911 |
| **TMEM16A** | 5OYG | Cryo-EM | digitonin | No | Mus musculus (mammal) | 260-266 |

| | | | | | | |
|---|---|---|---|---|---|---|
| **TMEM16A** | 5OYB | Cryo-EM | digitonin | 0.5mM $CaCl_2$ | Mus musculus (mammal) | 260-266,669-682 |
| TMEM16A | 5OYB | Cryo-EM + GROMACS or Amber/c36m | POPC | 2 $Ca^{2+}$ per chain, with 2 docked $PIP_2$ | Mus musculus (mammal) | 260-266,467-487,669-682 |
| TMEM16A | 7ZK3 – cluster 6 | Cryo-EM + GROMACS/c36m | POPC | 3 $Ca^{2+}$ per chain, with 1PBC removed | Mus musculus (mammal) | 260-266,467-482,526-527,669-682 |
| TMEM16A | 7ZK3 – cluster 8 | Cryo-EM + GROMACS/c36m | POPC | 3 $Ca^{2+}$ per chain, with 1PBC removed | Mus musculus (mammal) | 260-266,467-482,526-527,669-682 |
| TMEM16A | 7ZK3 – cluster 10 | Cryo-EM + GROMACS/c36m | POPC | 3 $Ca^{2+}$ per chain, with 1PBC removed | Mus musculus (mammal) | 260-266,467-482,526-527,669-682 |

***Appendix 1 - Table 2. Experimental structures selected for simulation over other similar structurers.***

| Selected Structure (PDB ID) | Similar Structure(s) (PDB ID) |
|---|---|
| 4WIS | 6QM9, 6QM5 |
| 6QMA | 6OY3 |
| 5OC9 | 6R65 |
| 7RXB | 7RXA, 7RX3, 6DZ7 |
| 7RXG | 7RX2, 7RWJ, 6E0H, 6E1O |
| 8B8J | 8B8M |
| 8TAG | 8SUR, 8SUN, 8TAI, 8TAL |
| 8BC0 | 8BC1 |
| 6P48 | 6P46, 6P49 |
| 5OYB | 7B5C, 7B5E |
| 5OYG | 7B5D |
